# Level Up the Brain! Novel PCA Method Reveals Key Neuroplastic Refinements in Action Video Gamers

**DOI:** 10.1101/2025.06.16.660043

**Authors:** Kyle Cahill, Mukesh Dhamala

**Affiliations:** Department of Physics and Astronomy, Georgia State University, Atlanta, GA, USA; Neuroscience Institute, Georgia State University, Atlanta, GA, USA; Center for Behavioral Neuroscience, Center for Diagnostics and Therapeutics, Georgia State University, Atlanta, GA, USA; Tri-Institutional Center for Translational Research in Neuroimaging and Data Science (TReNDS), Georgia State University, Georgia Institute of Technology, and Emory University, Atlanta, GA, USA

## Abstract

Action video games (AVGs) offer an ecologically rich experimental paradigm for studying how sustained cognitive demands drive behaviorally induced neuroplastic changes in the brain. We demonstrate that the neuroplastic refinements observed in long-term AVG players, referred to in this study as gamers, reflect more efficient neural mechanisms for reducing visuomotor information surprise during visuomotor decision-making by more effectively resolving internal conflict in competing motor plans, thus reducing uncertainty.

To explain how such adaptations unfold over time, we utilized the Cognitive Resource Reallocation (CRR) framework, defined as the dynamic redistribution of metabolic and functional resources to support behaviorally relevant neuroplastic adaptation under repeated, demanding task conditions. Using a novel region-cumulative principal component analysis (rcPCA) approach, we identified key brain regions that explain inter-subject variability, improving statistical power by isolating the most informative regions and reducing the burden of multiple comparisons.

Our findings suggest that prolonged AVG experience fosters more efficient visuomotor decision-making through top-down cognitive clarity, as reflected in the unobstructed transformation of learned value into goal-directed action, and bottom-up motor readiness, enabling improved visuomotor performance in gamers. These converging adaptations reduce internal conflict, mitigate uncertainty, and enable rapid yet skillful action selection. In sum, the brains of long-term gamers exhibit neuroplastic refinements consistent with CRR, marked by more effective transformation of sensory input into coherent motor output—an advantage especially critical in high-pressure environments. More broadly, these results illustrate how repeated cognitive challenge can perturb neurodynamic equilibria in ways that promote adaptive functional reorganization and enhanced cognitive ability.

## 1. Introduction

Video games provide a unique window into how complex, interactive environments shape the human brain. As one of the most widely consumed and cognitively engaging forms of entertainment globally^1^ video games have become a natural platform for studying experience-dependent neuroplasticity.^2, 3^ Different genres engage distinct cognitive processes from memory^4^ spatial navigation^5^ and problem-solving^6^, offering a spectrum of testable hypotheses about brain-behavior relationships. Among these, action video games (AVGs) stand out for their ability to consistently elicit high cognitive load^6^. They place players in fast-paced, sensory-rich environments that demand sustained attention, rapid visuomotor integration, and split-second decision-making.

Long-term exposure to AVGs has been associated with improved response times^7^, enhanced attentional control^8, 9^, more efficient sensorimotor learning^10^ and transformation ^11^, and cognitive flexibility^12^. These behavioral enhancements have even been shown to transfer to real-world domains such as surgery^13^, driving^14^, improving reading ability^15, 16^, and military training^17^. Yet, despite growing interest, it remains unclear how to organize the many observed neural mechanisms into unifying principles with necessary and sufficient explanatory power at the level of large-scale brain connectivity.^2, 3, 6, 18, 19^ Existing neuroimaging studies have reported changes in cortical thickness, gray matter volume,^5, 20, 21^ and white matter integrity in AVG players^22, 23^, referred to in this study as gamers, as well as altered activation in frontoparietal^15, 19^ and visual attention networks^16, 23, 24^. However, relatively few have integrated functional, structural, and directed connectivity within a single framework or tested whether these differences align with measurable behavioral outcomes.

In the context of action video game play, the “task” is not merely reacting to abstract stimuli or moving dots on a screen; it involves making rapid, high-stakes decisions to outmaneuver opponents, track fast-moving targets, and secure victory in dynamic, competitive matches. From the brain’s perspective, these actions occur within a meaningful social and performance-driven context, where success depends on real-time coordination and skillful execution. The brain is not optimizing to reduce reaction time as an end in itself; rather, it optimizes reaction time as a means to enhance overall performance, improving outcomes that are personally, socially, and competitively rewarding.

This study is grounded in the Cognitive Resource Reallocation (CRR) framework, which provides a systems-level explanation for how sustained cognitive demands reshape neural architecture^25^. CRR proposes that the brain optimizes performance by reallocating functional and metabolic resources toward anatomically plausible, behaviorally relevant circuits to more effectively meet strenuous task demands—ultimately striving to reduce prediction error^26, 27^ and thus enhancing behaviorally relevant neural processing. Rather than specifying the exact distribution of molecular mechanisms of meso-plasticity that give rise to brain-wide neuroplastic effects with behavioral relevance, which are altered in action video gamers^28^. CRR models neuroplasticity as a reallocation of resources toward circuits that support goal-directed behavior, focusing on neuroplastic changes within cognitive function (as defined by the APA)^29^ while moving away from less efficient processing.

In this study, our primary hypothesis is that long-term action video game play leads to neuroplastic reconfigurations that reduce visuomotor surprise—defined as unexpected visual or motor information during decision-making—through superior mechanisms for visuomotor uncertainty reduction and better handling of prediction compared to non-gamers. In tasks like our moving dots task, where the outcome is uncertain (e.g., a 50/50 chance), we suspect that the gamer’s brain does not predict a single outcome but rather anticipates both possibilities more readily. This ability to handle multiple potential outcomes demonstrates greater cognitive flexibility and attentional control, and cognitive features known to benefit action gamers^12, 30^. These adaptations would be reflected in connections that promote feedforward, goal-directed action selection and the integration of scene-specific visual contextual cues^31-34^, such as relative motion, which helps resolve uncertainty more efficiently. If confirmed, this would provide a necessary and sufficient explanation for the improved speed-accuracy tradeoffs observed in gamers, driven by the repeated need to make rapid, high-stakes decisions while playing an action video game.

Although we do not directly quantify visuomotor surprise, we aim to uncover signs of more effective methods of uncertainty reduction in action video game players compared to non-gamers during visuomotor decision-making. These signs include greater anticipatory/feedforward processing, motor readiness, and reduced internal conflict. Specifically, we expect gamers will show greater feedforward processing, motor readiness, improved anticipation, integration, and transformation of high-value visual cues into goal-directed action. This would reflect a more efficient, goal-directed neural organization when making visuomotor decisions. In contrast, non-gamers may rely more heavily on extended or compensatory uncertainty-reducing feedback loops, more resources towards early visual processing for object discrimination (e.g., of dot trajectories), and reward-seeking behavior that may be compensating for less effective goal-directed strategy and anticipatory action planning, indicating less efficient visuomotor transformations.

To test this hypothesis, we analyzed multimodal neuroimaging data from gamers and non-gamers who participated in a modified moving-dots task designed to probe visuomotor decision-making^7^. The results revealed that long-term action video gamers exhibited faster response times (∼190 ms) with marginal gains in accuracy (∼2-3%) compared to non-gamers. In decision-making literature, speed-accuracy trade-offs are typically accepted as the norm^35, 36^. Therefore, this unexplained effect requires a necessary and sufficient explanation. We analyzed structural connectivity (SC) using thirteen different SC measures provided by DSI Studio^37^, functional connectivity (FC), and directed time-domain Granger causality (dFC) in this study. During PCA analysis, we extracted submatrices of the dFC data into ‘sender’ (outflow) and ‘receiver’ (inflow) contributions, reflecting the directionality of influence between brain regions. The ‘total’ dFC influence represents the combined interaction between these regions, incorporating both sender and receiver contributions. In addition to undirected and directed connectivity analyses, we examined combined structure-function measures including SFC undirected coupling and SdFC sender-mode coupling to explore alignment between a region’s structural substrate and its capacity for equivalent functional load, such as undirected synchrony (SFC) or outbound influence (SdFC sender).

Whole-brain connectivity matrices are high-dimensional, comprising tens of thousands of pairwise connections, which posed significant challenges. Initial attempts to isolate meaningful whole-brain functional effects using conventional statistical thresholding yielded an overwhelming number of results across the AAL3 atlas’ 166 × 166 pairwise connections^38^, with no single principled method to distinguish signal from noise. Assessing all directed connections among 166 regions of the AAL3 atlas yields over 27,000 comparisons in anatomically unconstrained dFC data and over 13,000 comparisons in anatomically unconstrained symmetric FC data, despite a modest sample of 47 recruited participants, which severely diminished statistical power.

To address this challenge, a complementary data-driven PCA-based approach ^39^ was developed to sweep across the entire dataset and identify the strongest sources of inter-subject signal variance. Unlike traditional PCA approaches that reduce time series or select top ROIs from early components, the region-cumulative PCA (rcPCA) method utilizes cumulative variance-weighted contribution scores across all components, up to a defined cumulative explained variance threshold. This balances the goal of capturing meaningful inter-subject variation while excluding spurious components that reflect noise or isolated variance.

An ICA-based method to dimensionality reduction was initially considered, but rcPCA was ultimately developed as a more suitable alternative for this study’s goal of investigating how AVG experience shapes SC, FC, dFC, SFC, and SdFC (sender) profiles within a predefined anatomical atlas such as AAL3. Independent Component Analysis (ICA) is a powerful tool for uncovering distributed functional networks via blind source separation, using higher-order statistics such as kurtosis or negentropy, and has yielded valuable insights into large-scale brain organization ^40, 41^. While ICA excels at detecting latent sources and identifying network-level structure, it does not natively support anatomically localized, region-level ranking. Techniques such as spatial template matching, dual regression, ICN labeling, and joint ICA each extend ICA’s utility, with template matching aiding anatomical alignment, dual regression estimating subject-level component expression, ICN labeling providing network-level categorization, and joint ICA enabling multimodal data fusion. However, none of these approaches cleanly resolve the core challenge of identifying which anatomical regions within a predefined structural atlas like AAL3 contribute most meaningfully to inter-subject variability across connectivity modalities.

We propose, as our second hypothesis, that the rcPCA method will identify behaviorally relevant connections and structure–function couplings, offering a richer understanding of the neuroplastic adaptations associated with long-term action video game play. This, in turn, provides the means to rigorously examine our primary hypothesis that the behavioral advantage in visuomotor decision-making observed in gamers plausibly stems from a cognitive resource reallocation away from regions and connections involved in uncertainty-reducing circuits during visuomotor decisions, and reflects a reduction in visuomotor surprise.

## 2. Results

Our ROI selection strategy followed standard PCA-based practices^42, 43^ commonly used in neuroimaging studies^39, 43^, prioritizing high-loading features from each principal component ^44^. Specifically, we selected the top 20 contributing ROIs per component to emphasize regions that most strongly explained structured variance while minimizing the inclusion of low-weight contributors. This approach aligns with established PCA interpretation techniques, which typically focus on dominant features from early components to maximize interpretability.

As shown in Figure 1, PCA-based ROI contributions exhibited strong correspondence with raw variance rankings across multiple brain connectivity metrics, including functional connectivity (FC), directed sender connectivity, and total Granger causality. Each modality yielded Spearman correlations exceeding 0.85 across the top-ranked ROIs, with significance (p < 0.001) occurring rapidly as sample size (n) increased (see Figure 2,b). Although FC showed slightly weaker monotonic rank alignment compared to directed connectivity, it still displayed statistically robust convergence by n = 16, indicating reliable structure in the PCA-derived regional variance. Validation of rcPCA-derived ROIs utilized a combination of rank correlation, permutation testing, and hypergeometric overlap statistics. Using a significance threshold of p < 0.001, this approach confirmed that the observed alignments between PCA-derived ROI rankings and raw variance were statistically robust and highly unlikely to occur by chance.

**Figure 1.**
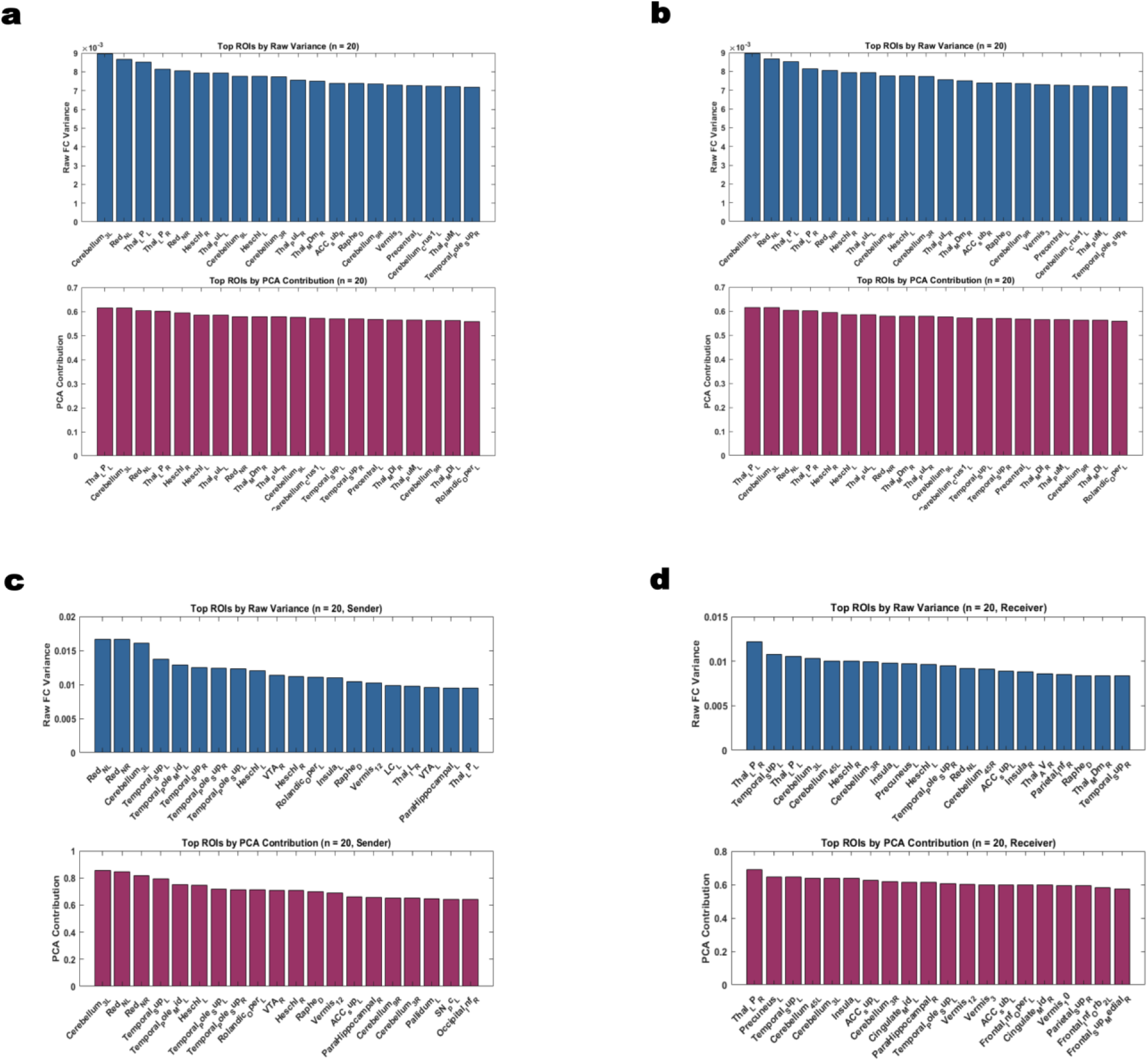
Top 20 PCA Selected ROIs from FC and dFC Based on Explained Variance 80%. Principal component analysis (PCA) was applied to two functional connectivity matrices: undirected functional connectivity (FC) and directed functional connectivity (dFC). The dFC matrix was further analyzed in three forms: total (collapsed directionality), sender (row-wise), and receiver (column-wise). For each, ROIs were ranked by their contribution to across-subject variance, and the top 20 ROIs were selected based on a cumulative 80% explained variance threshold. **(a) FC:** undirected Pearson correlation matrix. **(b) dFC (total):** Granger causality values summed across source and target roles to capture total directed influence. **(c) dFC (sender**): variance contributions based on a region’s outgoing directed influences. **(d) dFC (receiver**): variance contributions based on a region’s incoming directed influences. This decomposition enables the identification of ROIs with asymmetric functional dynamics and highlights regions that dominate information transmission versus reception.

**Figure 2.**
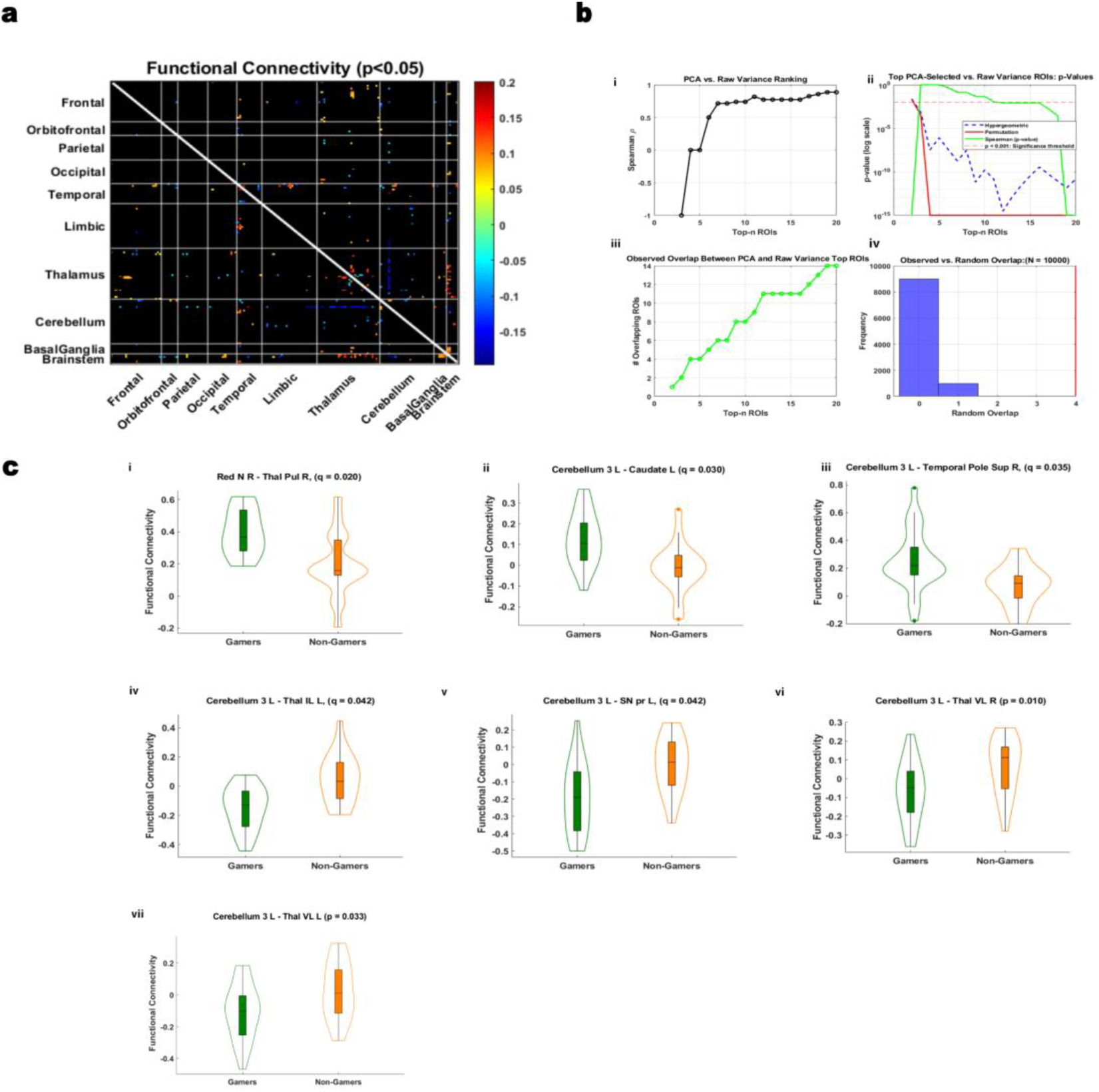
Functional Connectivity Differences Involving Top rcPCA ROIs. Multi-step analysis of functional connectivity (FC) group differences and PCA-based ROI selection. **(a)** Whole-brain FC matrix showing significant group differences (p < 0.05, uncorrected), organized by anatomical region. **(b)** Validation of the PCA-based method: (i) Spearman’s rank correlation (ρ) across top-n selections comparing PCA-weighted and raw variance-based rankings; (ii) Comparison of statistical sensitivity using Spearman correlation, hypergeometric overlap p-values, and permutation-derived p-values between PCA- and raw-ranked ROIs; (iii) Overlap between PCA- and raw-ranked ROIs increases systematically with top-n selections; (iv) PCA-selected ROI overlap exceeds chance across 10,000 permutations. **(c)** Violin plots illustrating significant FC differences (q < 0.05, FDR-corrected) involving top PCA-selected ROIs in cerebellar, subcortical, midbrain, thalamic, and temporal areas

### 2.1 Group Differences in Functional and Directed Connectivity

#### 2.1.1 Group Differences in Functional Connectivity

Whole-brain FC analysis revealed significant group differences (p < 0.05) between gamers and non-gamers, concentrated in thalamic, cerebellar, midbrain, and temporal regions (Figure 2a). PCA-selected ROIs, validated in Figure 2b, showed robust effects following FDR correction. Gamers exhibited significantly stronger functional connectivity between the right red nucleus and the right pulvinar thalamic nucleus (*p* = 0.0061, *q* = 0.02, *d* = 0.85), and between left cerebellar lobule 3 and both the right superior temporal pole (*p* = 0.0014, *q* = 0.035, *d* = 1.12) and the left caudate nucleus (*p* = 0.00093, *q* = 0.03, *d* = 1.00). In contrast, non-gamers showed stronger connectivity between left cerebellar lobule 3 and the left intralaminar thalamic nuclei (*p* = 0.00085, *q* = 0.042, *d* = –1.26), the left substantia nigra pars reticulata (*p* = 0.002, *q* = 0.042, *d* = –1.10), and both the right and left ventrolateral thalamic nuclei (*p* = 0.01, *d* = –0.76; *p* = 0.03, *d* = –0.87, respectively). These effects reflect consistent, large-magnitude differences in functional connectivity across subcortical–cerebellar circuits. Absolute effect sizes were consistently large, ranging from *d* = 0.76 to 1.26 across significant connections.

Full statistical results, including all p-values, FDR-adjusted q-values, and Cohen’s d effect sizes for each comparison, are reported in Supplementary Figure 2.

#### 2.1.2 Group Differences in Directed Functional Connectivity

Directed connectivity analyses using Granger causality revealed significant differences in directed functional connectivity (dFC) between gamers and non-gamers across sender, receiver, and total modes (*p* < 0.05, uncorrected). These effects were concentrated in midbrain, thalamic, cerebellar, and anterior cingulate cortex (ACC) regions (Figure 3a–c). In sender mode, non-gamers exhibited stronger directed influence from the left supracallosal ACC to the right ventrolateral thalamus (*p* = 0.0003, *q* = 0.042, *d* = –0.91), the only connection that survived false discovery rate (FDR) correction. Additional uncorrected differences included stronger non-gamer outflow from the left ACC to the left cerebellar lobule 7b (*p* = 0.004, *d* = –0.65) and from the right ventral tegmental area (VTA) to both the right ventrolateral thalamus (*p* = 0.006, *d* = –0.96) and the left cerebellar lobule 3 (*p* = 0.012, *d* = –0.75).

**Figure 3.**
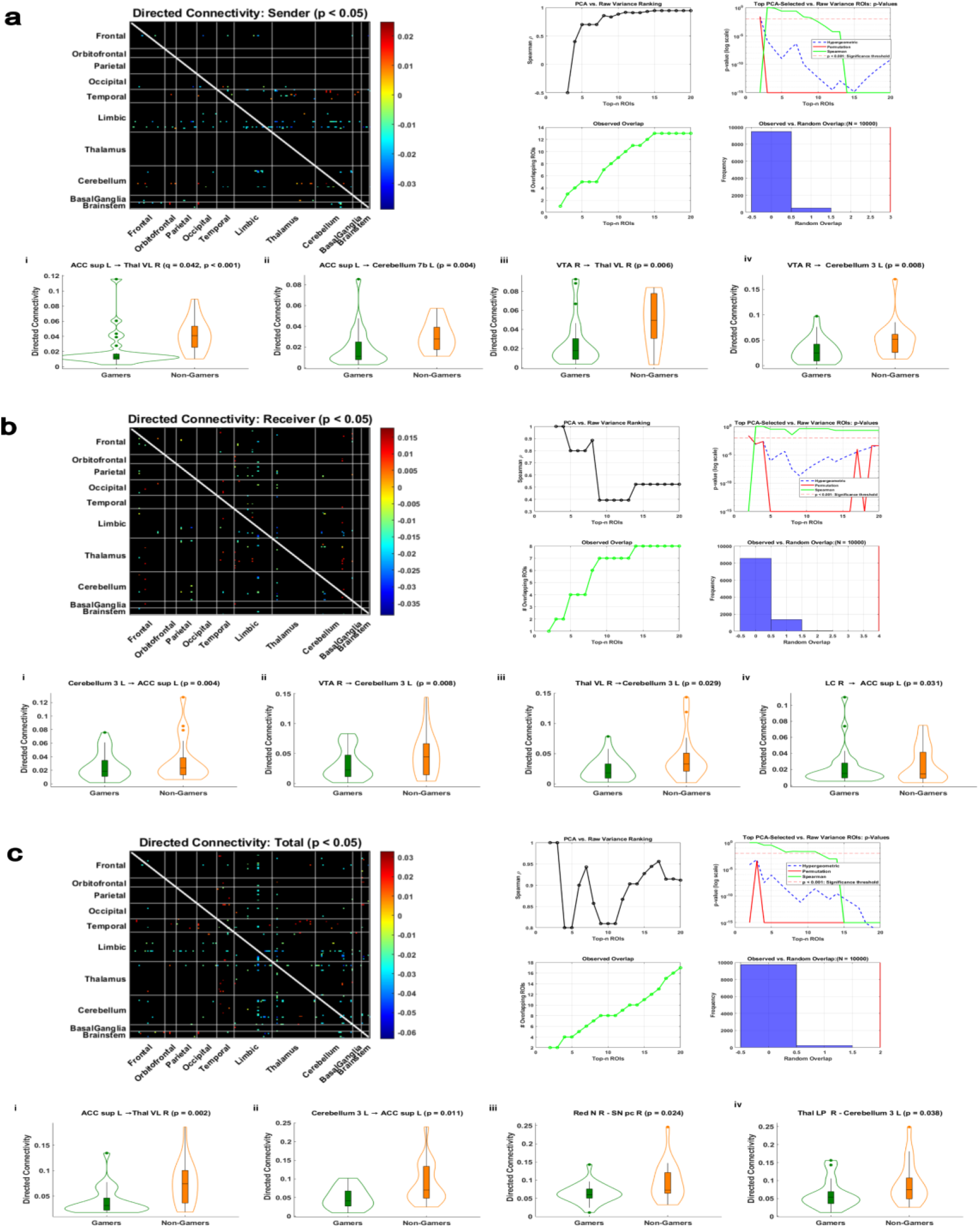
Directed Connectivity Differences Involving Top rcPCA ROIs. Group-level differences in directed functional connectivity (dFC) between gamers and non-gamers using PCA-based ROI selection. Each panel includes a full dFC matrix of significant group differences (p < 0.05), validation of PCA-based selection stability, and violin plots highlighting the strongest observed effects (p < 0.05). **(a)** dFC (sender): group differences in outgoing influence. A connection from the anterior cingulate cortex (ACC sup L) to the right thalamus (VL R) survived FDR correction (q < 0.05). Additional uncorrected effects were observed from VTA R to cerebellar and thalamic targets. **(b)** dFC (receiver): group differences in incoming influence. Effects were observed in Cerebellum 3 L receiving projections from VTA R and Thal VL R, and in ACC sup L receiving input from LC R.**(c)** dFC (total): group differences in direction-collapsed influence (sum of sender and receiver roles). Notable effects included those involving ACC sup L, SN pc R, and Cerebellum 3 L.

In receiver mode, non-gamers showed greater inflow to the left supracallosal ACC from the left cerebellar lobule 3 (*p* = 0.004, *d* = –1.00) and the right locus coeruleus (*p* = 0.031, *d* = –0.83), while bilateral VTA and right thalamus also showed elevated input to the left cerebellum. Total mode effects reflected overlapping but distinct connectivity patterns, including stronger non-gamer influence from the left ACC to the right thalamus (*p* = 0.002, *d* = –0.98) and from the left cerebellar lobule 3 to the left ACC (*p* = 0.011, *d* = –0.87). Effect size magnitudes across modes ranged from *d* = 0.59 to 1.00, indicating robust directional asymmetries in dFC across groupsFull statistical results, including all p-values, FDR-adjusted q-values, and Cohen’s d effect sizes for each comparison, are displayed in Supplementary Figure 2.

### 2.2. Extensions of PCA-Based ROI Selection Beyond Functional Connectivity

#### 2.2.1 Group Differences in Structural Connectivity

Structural connectivity (SC) analysis using diffusion measures, fractional anisotropy (FA), axial diffusivity (AD), isotropy (ISO), and non-restricted diffusion imaging (NDRI) revealed significant group-level differences between gamers and non-gamers involving rcPCA-derived ROIs (*p* < 0.05), as shown in Figure 4. For FA, reduced connectivity in non-gamers was observed between the left calcarine cortex and the left superior occipital gyrus (*p* = 0.036, *d* = –0.78), a connection also associated with slower response times in behavioral analysis (see Figure 6).

**Figure 4.**
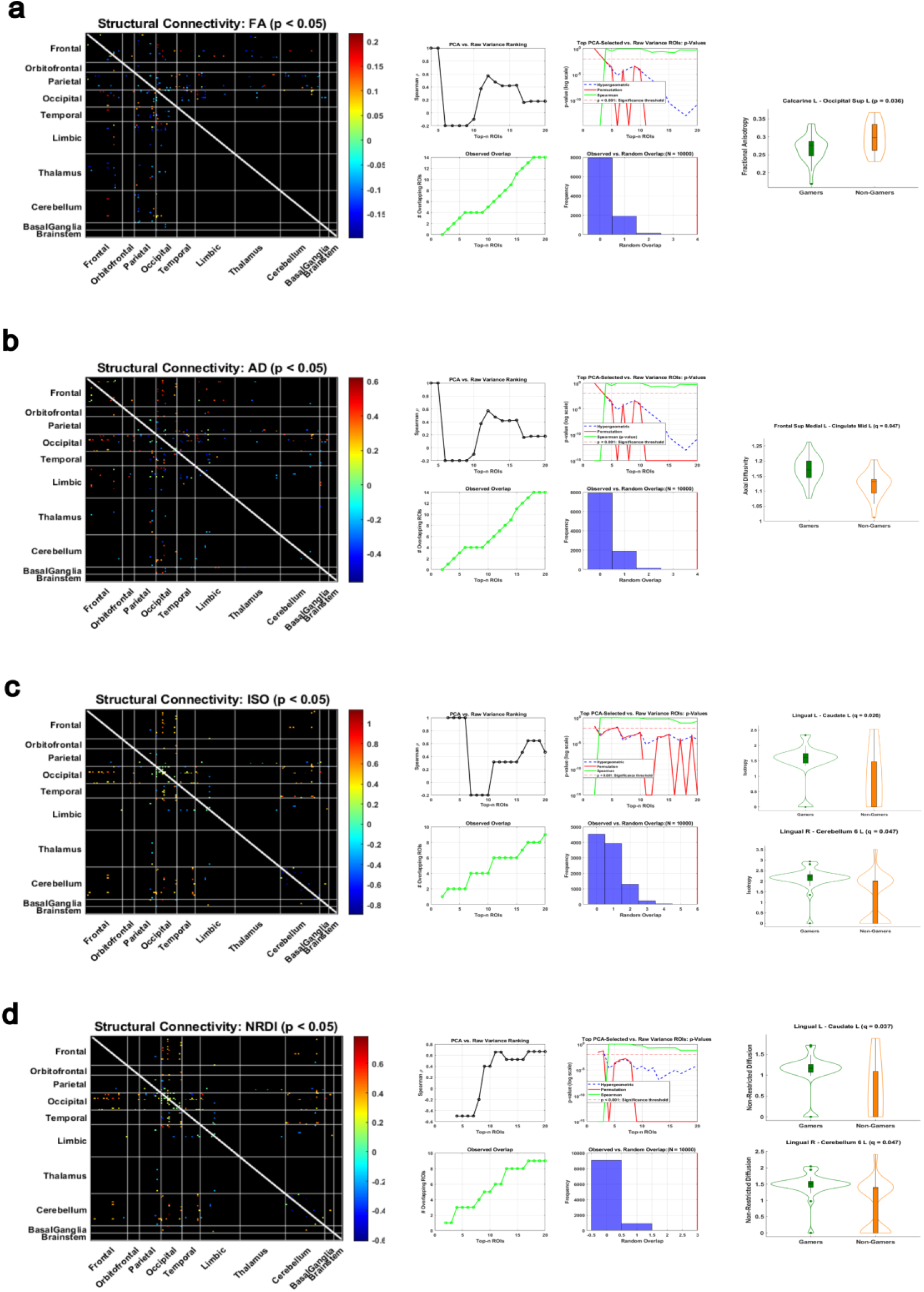
Structural Connectivity Differences Filtered by Top rcPCA ROIs. Group-level differences in structural connectivity (SC) between gamers and non-gamers using PCA-based ROI selection across four microstructural diffusion metrics. Each panel shows a full SC matrix of significant group differences (p < 0.05), validation of PCA-based selection stability, and violin plots of the strongest observed effects (p < 0.05). PCA-based filtering revealed consistent and interpretable differences across SC measures. All effects shown reflect comparisons involving top PCA-selected ROIs **(a)** Fractional anisotropy (FA): a significant difference was observed in the connection between Calcarine L and Occipital Sup L, which was also associated with slower response times in behavioral analysis. **(b)** Axial diffusivity (AD): Group differences were observed in connections involving Frontal Sup Medial L and Cingulate Mid L. **(c)** Isotropic volume fraction (ISO): effects were found in Lingual L – Cerebellum 6 L and Lingual R - Cerebellum 6 L. **(d)** Non-restricted diffusion imaging (NRDI): the same cerebellar projections from Lingual L and Lingual R remained significant.

In the AD measure, a robust group difference emerged between the left superior medial frontal gyrus and the left middle cingulate cortex (*p* = 0.0003, *q* = 0.047, *d* = 1.16), the only FA or AD comparison to survive FDR correction. ISO-based analyses revealed stronger connectivity in gamers between the left lingual gyrus and the left caudate (*p* = 0.0002, *q* = 0.026, *d* = 1.22) and between the right lingual gyrus and the left cerebellar lobule 6 (*p* = 0.0007, *q* = 0.047, *d* = 0.96). These region pairs also showed significant differences in the NDRI measure, with both comparisons maintaining significance following FDR correction.

Absolute effect sizes across all measures were consistently large, with *d* values ranging from 0.78 to 1.22. Full statistical results, including all p-values, FDR-adjusted q-values, and Cohen’s d effect sizes for each comparison, are reported in Supplementary Figure 2.

#### 2.2.2 Structure–Function Coupling

Structure-function coupling measures the alignment between a region’s structural connectivity measure and its corresponding capacity for functional load^45^. To quantify this relationship, Pearson correlations were computed between structural connectivity (SC) and both functional connectivity (FC) and directed functional connectivity (dFC sender mode). The undirected correlation with FC is referred to as SFC, while the correlation with sender-mode dFC is termed SdFC (sender). As shown in Figure 5a, gamers exhibited significantly stronger coupling in the cerebellum. These included Vermis 3 with mean diffusivity (*p* = 0.046, *d* = 0.64), Vermis 9 with both mean length (*p* = 0.024, *d* = 0.64) and fractional anisotropy (*p* = 0.049, *d* = 0.55), and Cerebellum 10L with fractional anisotropy (*p* = 0.046, *d* = 0.65) and quantitative anisotropy (*p* = 0.049, *d* = 0.65).

**Figure 5.**
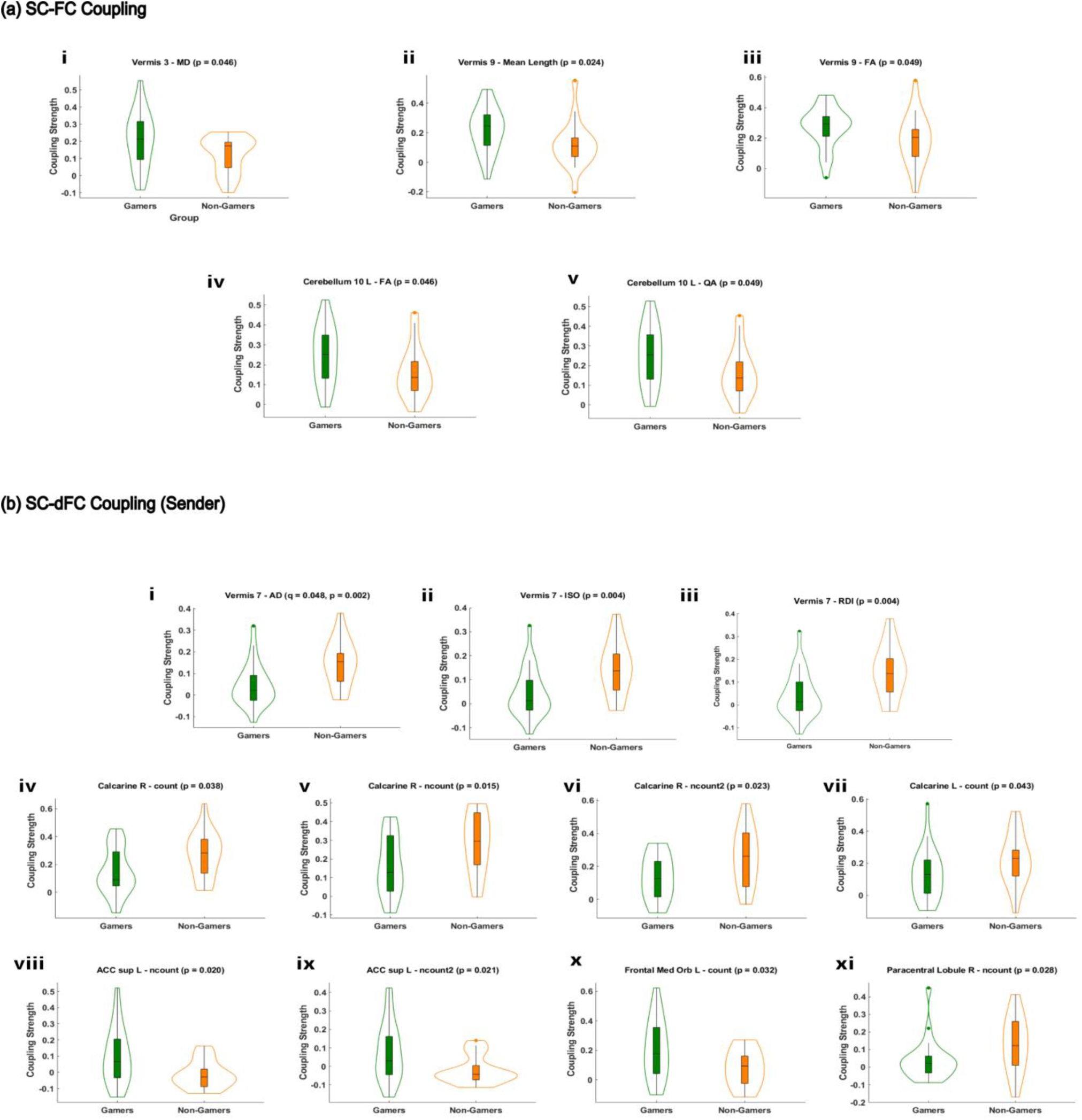
SC-FC and SC-dFC (Sender) Coupling Differences Involving Top rcPCA ROIs. Group-level differences in structure–function coupling between gamers and non-gamers, using PCA-based ROI selection. **(a)** SC–FC coupling: Significant group differences (p < 0.05) were observed across several structural connectivity measures, including mean diffusivity (MD), mean length, fractional anisotropy (FA), and quantitative anisotropy (QA), in connections involving Vermis 3, Vermis 9, and Cerebellum 10 L.**(b)** SC–dFC coupling (sender): Group differences in sender-based structure–function coupling strength (p < 0.05), with significant effects observed in Vermis 7 across intensive measures such as axial diffusivity (AD), isotropy (ISO), and restricted diffusion imaging (RDI). Significant effects were also observed within the Calcarine cortex (R and L), ACC sup L, Frontal Med Orb L, and Paracentral Lobule R when coupled with extensive measures such as count, ncount, and ncount2. All effects reflect comparisons involving top PCA-selected ROIs. Sender-based SC–dFC coupling was selected to parallel SC–FC coupling based on similar FC and dFC PCA validation profiles prior to coupling. Validation of PCA-based ROI selection followed the same procedure as in previous analyses of FC, dFC, and SC connectivity.

In the SdFC sender condition shown in Figure 5b, non-gamers showed significantly stronger coupling in Vermis 7 across multiple measures, including axial diffusivity (*p* = 0.002, *q* = 0.048, *d* = –1.00), isotropy (*p* = 0.004, *d* = –0.97), and restricted diffusion (*p* = 0.004, *d* = –0.97). Additional effects favoring non-gamers were found in the right calcarine cortex for count (*p* = 0.038, *d* = –0.74), ncount (*p* = 0.015, *d* = –0.82), and ncount2 (*p* = 0.023, *d* = –0.82), as well as in the left calcarine cortex (*p* = 0.043, *d* = –0.59).Gamers, in contrast, exhibited stronger SdFC sender coupling in the left supracallosal anterior cingulate cortex for both ncount (*p* = 0.020, *d* = 0.82) and ncount2 (*p* = 0.021, *d* = 0.83), in the left medial orbitofrontal cortex for count (*p* = 0.032, *d* = 0.74), and in the right paracentral lobule for ncount (*p* = 0.028, *d* = –0.67).

Effect size magnitudes were moderate to large across both coupling modes, with *d* values ranging from 0.55 to 1.00. The Vermis 7–axial diffusivity pair was the only comparison to survive FDR correction. These findings highlight distinct structure–function coupling profiles between groups, particularly in cerebellar and occipital regions. Full statistical results, including all p-values, FDR-adjusted q-values, and Cohen’s d effect sizes for each comparison, are reported in Supplementary Figure 2.

### 2.3 Brain–Behavior Relationships

Response time was significantly associated with connectivity strength and structure–function coupling across modalities. Faster responses were linked to negatively sloped Spearman correlations, while slower responses were associated with positively sloped trends.

In functional connectivity, faster responses were linked to stronger connectivity between the right red nucleus and the right pulvinar thalamus (*r* = –0.41, *p* = 0.009), whereas slower responses were associated with increased connectivity in cerebellar and thalamic circuits, including the left cerebellar lobule 3 and the right substantia nigra pars compacta (*r* = 0.33, *p* = 0.036), as well as the left ventrolateral thalamus and the left cerebellar lobule 3 (*r* = 0.33, *p* = 0.037).

In directed connectivity, slower responses were correlated with increased influence from the right ventral tegmental area to the left cerebellar lobule 3 (*r* = 0.41, *p* = 0.008), from the right substantia nigra pars compacta to the right red nucleus (*r* = 0.40, *p* = 0.003), and from the left supracallosal anterior cingulate cortex to the left cerebellar lobule 7b (*r* = 0.34, *p* = 0.032). Additional significant effects included connections from the right lateral posterior thalamus to the left cerebellar lobule 3 (*r* = 0.40, *p* = 0.009) and from the right locus coeruleus to the left supracallosal anterior cingulate cortex (*r* = 0.40, *p* = 0.010).

In structural connectivity, slower responses were associated with higher fractional anisotropy between the left calcarine cortex and the left superior occipital gyrus (*r* = 0.35, *p* = 0.020), while faster responses were linked to lower fractional anisotropy between the left parahippocampal gyrus and the left precuneus (*r* = –0.36, *p* = 0.011), and to lower quantitative anisotropy between the left superior temporal pole and the left orbital part of the inferior frontal gyrus (*r* = –0.42, *p* = 0.005).

In structure–function coupling, faster response times were associated with stronger coupling in the right mid-occipital cortex, including axial diffusivity (*r* = –0.31, *p* = 0.050) and non-restricted diffusion imaging (*r* = – 0.32, *p* = 0.042). Stronger coupling in the right supracallosal anterior cingulate cortex was also associated with faster responses across multiple measures: axial diffusivity (*r* = –0.44, *p* = 0.006), fractional anisotropy (*r* = – 0.44, *p* = 0.006), normalized quantitative anisotropy *(r* = –0.39, *p* = 0.015), and non-restricted diffusion imaging (r = –0.37, *p* = 0.020). In sender-mode coupling, faster responses were further linked to greater structure–function alignment from the right anterior cingulate cortex (*e.g.,* FA: *r* = –0.44, *p* = 0.006; NRDI: *r* = –0.37, *p* = 0.020; NQA: *r* = –0.39, *p* = 0.015), as well as from the left medial orbital frontal cortex (*r* = –0.34, *p* = 0.035) and the left calcarine cortex (*r* = –0.34, *p* = 0.032). All brain–behavior correlations supporting these effects are displayed in Figure 6.

**Figure 6.**
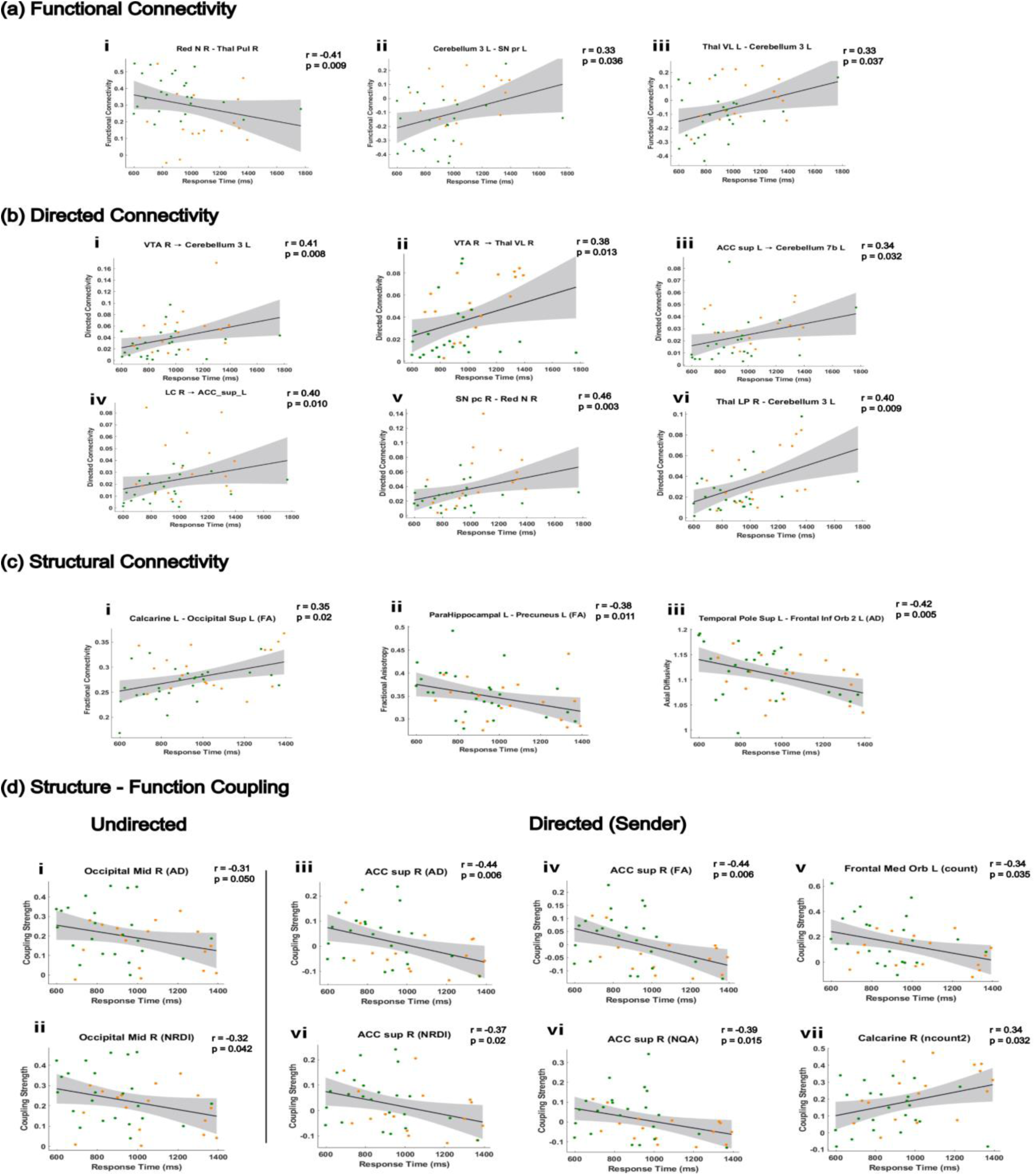
Behaviorally Relevant Connectivity and Coupling Involving Top PCA ROIs. Brain–behavior relationships linking connectivity strength to response time across functional, structural, and structure–function coupling modalities. Positive correlations indicate slower responses; negative correlations indicate faster responses. **a**) Functional connectivity (FC): Faster response times were associated with stronger FC between the red nucleus and thalamus (PuL R). In contrast, slower responses were associated with stronger FC between midbrain (SN pr L) and thalamic (Thal VL L) regions connected to the cerebellum. **(b)** Directed connectivity (dFC): Slower responses were linked to increased directed influence among midbrain (VTA R, SN pc R), thalamic (Thal VL R), cerebellar, and anterior cingulate (ACC sup L) pathways. **(c)** Structural connectivity (SC): FA and AD-based metrics showed significant associations with behavior. Higher FA between Calcarine L and Occipital Sup L predicted slower performance **(d)** Structure–function coupling: Both SC–FC and sender-based SC–dFC coupling strength tracked with response time. Faster responses were associated with stronger coupling in the frontal, anterior cingulate, and mid-occipital regions.

## 3. Discussion

### 3.1 A Novel PCA-Based Framework for Region Selection

To address the challenges of navigating the complexity of whole-brain connectivity matrices, we developed a novel PCA-based framework that reimagines the role of principal component analysis (PCA) in neuroimaging. Rather than relying on anatomical constraints or arbitrary statistical cutoffs, this method decomposes subject-by-connection matrices into orthogonal principal components and computes the eigenvalue-weighted sum of absolute contributions for the top user-defined regions across all components. The result is a global variance contribution score for each brain region, allowing us to prioritize the most informative regions for downstream group comparisons based on their contribution to explained variance. For our analysis, we selected the top 20 regions across all components, up to an explained variance threshold of 80%, to emphasize the most relevant contributors for downstream analysis while controlling for stability.

Crucially, this region-cumulative PCA method (rcPCA) leverages the mathematical structure of PCA^39, 46^ to form an orthonormal basis over the space of brain regions, enabling a principled and interpretable ranking of ROI importance. By weighting regional contributions according to the variance explained by each principal component, the method prioritizes dominant, structured inter-subject variance. This orthogonal decomposition avoids redundancy, permitting straightforward accumulation of variance-weighted contributions.

Moreover, rcPCA flexibly handles both undirected and directed connectivity matrices. For directed measures, such as Granger causality, the method separately calculates contribution scores for the sender, receiver, and total contributions, allowing for a nuanced interpretation of directional asymmetries in brain networks. This increases the interpretability and relevance of the selected ROIs. Altogether, this novel application of PCA provides a mathematically robust alternative to traditional thresholding and anatomical filtering, focusing on variance-based selection that generalizes across structural, functional, directed connectivity, and coupling modalities.

We developed a multi-pronged validation strategy to evaluate the internal consistency and robustness of rcPCA using rank correlation, permutation testing, and hypergeometric overlap statistics with a conservative significance threshold of p < 0.001 as our primary validation criteria. This strategy enabled a drastic reduction in the number of statistical comparisons required for downstream testing, significantly improving our ability to detect FDR-corrected (q < 0.05) results that would have been obscured by multiple comparison corrections.

In our previous study, we applied a structurally constrained functional neuroimaging analysis, limiting connectivity assessments to anatomically plausible tracts informed by white matter tractography^25^. While this approach enhanced biological plausibility, it was inherently limited by tractography reconstruction assumptions and macro-level user-defined parameters such as allowable streamline length and angular thresholds. The rcPCA method offers a complementary perspective by identifying dominant, behaviorally relevant patterns of variance without relying on anatomical priors. Unlike anatomical filtering, which can miss regions not tightly bound to known white matter pathways, PCA captures emergent structure across subjects and modalities. By combining tractography-informed constraints to ensure biological plausibility with rcPCA to filter for the most robust patterns in inter-subject variability, we constructed a methodological framework capable of supporting a comprehensive and principled investigation of the neuroplastic adaptations associated with video game experience.

### 3.2 Group Differences Between Gamers and Non-Gamers Across Modalities

Group-level comparisons between gamers and non-gamers revealed widespread differences across multiple connectivity domains, all of which involved at least one PCA-identified region of interest (ROI).

In functional connectivity (FC), shown in Figure 2, gamers exhibited significantly stronger FC between the right red nucleus and the right pulvinar thalamus, the left cerebellum lobule 3 and the left caudate nucleus, and the left cerebellum lobule 3 and the right superior temporal pole. In contrast, non-gamers showed stronger FC between the left cerebellum lobule 3 and the left intralaminar thalamus, the left cerebellum lobule 3 and the left substantia nigra pars reticulata, and the left cerebellum lobule 3 and the ventrolateral thalamus bilaterally. All FC differences, except for the connections involving the ventrolateral thalamus, survived FDR correction.

Several of these FC connections were also significantly correlated with response time (RT), indicating behavioral relevance. These patterns suggest that non-gamers may rely more heavily on feedback-based loops involving cerebellar–thalamic–midbrain circuits, potentially as a mechanism to reduce uncertainty during motor action selection. Effect sizes were consistently large, with Cohen’s *d* magnitudes ranging from 0.76 to 1.26.

In directed functional connectivity (dFC), presented in Figure 3, sender-mode results revealed a robust group difference in directed influence favoring non-gamers from the left superior anterior cingulate cortex to the right ventrolateral thalamus, which survived FDR correction. Additional uncorrected differences favoring non-gamers included projections from the left superior anterior cingulate cortex to the left cerebellum lobule 7b, from the right ventral tegmental area to the right ventrolateral thalamus, and from the right ventral tegmental area to the left cerebellum lobule 3. Notably, each of these connections also showed significant positive correlations with RT.

Receiver-mode differences also favored non-gamers, with several directed connections showing stronger influence toward key target regions. These included projections from the left cerebellum lobule 3 to the left superior anterior cingulate cortex, from the right ventral tegmental area to the left cerebellum lobule 3, and from the right locus coeruleus to the left superior anterior cingulate cortex. Several of these connections also showed significant positive correlations with RT. Total-mode results closely mirrored the sender- and receiver-mode patterns, with notable unique group differences such as increased connectivity from the right red nucleus to the right substantia nigra pars compacta and from the left thalamus to the left cerebellum lobule 3 in non-gamers.

The consistency of these effects across all directed functional connectivity (dFC) modes, coupled with moderate to large effect sizes (*d* = 0.59 to 1.00), reinforces the earlier findings from tractography-constrained analyses^25^, which revealed that non-gamers exhibit a greater number of directed connections. The current results build on that observation, suggesting that non-gamers engage in a broader, more distributed pattern of targeted information flow, likely reflecting compensatory network recruitment to signal the need for an imminent motor response during rapid visuomotor decision-making.

In structural connectivity (SC) analysis using diffusion measures (FA, AD, ISO, and RDI), we observed group-level differences involving PCA-derived regions of interest (*p* < 0.05) as shown in Figure 4. For FA, a notable and seemingly counterintuitive difference emerged that FA was significantly higher in non-gamers between the left calcarine cortex and the left superior occipital gyrus—early visual areas^47^, which also had a positive correlation with RT thus associated with slower response times in behavioral analysis (see Figure 6). This finding suggests that non-gamers may over-rely on early-stage visual processing to compensate for inefficiencies further downstream. For AD, group differences emerged between the left superior medial frontal gyrus and the left mid-cingulate cortex (*q* < 0.05). ISO revealed significant effects in connections between the right lingual gyrus and the left cerebellum lobule 6, as well as between the left lingual gyrus and the left caudate nucleus. Both connections remained significant in NRDI and under FDR correction. Absolute effect sizes across all SC measures were substantial, with Cohen’s *d* magnitudes ranging from 0.78 to 1.22, indicating consistently large group-level differences.

Structure–function coupling findings displayed in Figure 5 provided further insight into group differences by measuring the alignment between a region’s structural connectivity profile and its capacity for functional load^45^. Gamers exhibited stronger structure–function coupling (SFC) within cerebellar regions, including vermis lobule 3 (MD), vermis lobule 9 (mean length, FA), and the left cerebellum lobule 10 (FA, QA). In contrast, non-gamers showed stronger structure–directed functional coupling (SdFC sender) in bilateral calcarine cortices and vermis lobule 7. One of these connections, involving vermis lobule 7 and AD-based SdFC sender coupling, survived FDR correction.

Gamers also demonstrated stronger SdFC sender coupling in key frontal and cingulate regions, including the left superior anterior cingulate cortex (ncount, ncount2), the left medial orbital frontal cortex (count), and the left paracentral lobule (ncount). These findings underscore a functional distinction between undirected synchrony, as measured by SFC, and directional signaling capacity, as captured by SdFC sender. Stronger coupling in gamers may reflect more efficient transmission channels for dynamic signaling, consistent with prior evidence of increased dorsal attention to salience network (DAN-to-SN) switching, and which the superior anterior cingulate cortex as a core node in the salience network (SN) ^24^.

### 3.3 Brain–Behavior Relationships

Connectivity and coupling strengths across all modalities were significantly associated with response time (RT) (Figure 6). These associations reinforce the behavioral relevance of group differences and suggest that the specific network configurations observed in gamers versus non-gamers map divergent visuomotor strategies.

Non-gamers exhibited more engagement in regions and connections involving cerebellar–midbrain–thalamic circuits, which were consistently associated with slower responses and may reflect a less efficient allocation of cognitive resources.

#### 3.3.1 Functional Connectivity (FC)

Faster RTs, indicated by negative correlations, were linked to stronger FC between the red nucleus and thalamus pulvinar. In contrast, slower RTs were associated with stronger FC in cerebellar-thalamic/cerebellar-midbrain, connections particularly between cerebellum 3L and thalamus intralaminar L, substantia nigra pars reticulata L, and thalamus ventrolateral L/R—all of which favored non-gamers. These patterns suggest that overreliance on cerebellar-thalamic error correction signals ^48^, which would slow down visuomotor transformation, promote tighter inhibitory control, evidenced by increased substantia nigra-thalamus ventrolateral synchrony in non-gamers.^49^ This elevated substantia nigra–thalamus ventrolateral synchrony suggests reduced motor readiness, consistent with increased inhibitory gating of thalamic output, and reflects a diminished capacity to swiftly resolve competing motor plans^50^ ultimately delaying commitment to action selection under uncertainty and prolonging response time.

#### 3.3.2 Directed Functional Connectivity (dFC)

All three sender-mode connections that favored non-gamers—anterior cingulate cortex superior L → cerebellum 7b L, ventral tegmental area R → thalamus ventrolateral R, and ventral tegmental area R → cerebellum 3L—also showed significant positive correlations with RT, indicating slower responses. Likewise, several receiver-mode connections that favored non-gamers (e.g., cerebellum 3L → anterior cingulate cortex superior L, ventral tegmental area R → cerebellum 3L, locus coeruleus R → anterior cingulate cortex superior L) were also positively correlated with RT. These results point to a more distributed routing of information in non-gamers associated with cerebellar error correction^51^, stress response^52^ (locus coeruleus), and dopaminergic reward-seeking impulsiveness^53^ (ventral tegmental area) VTA. In this context, this increased signaling from VTA may be acting as a compensatory mechanism for a lack of goal-directed strategy, one incurs a performance cost.

#### 3.3.3 Structural Connectivity (SC)

Regarding structural connections, the FA connection between the calcarine cortex and superior occipital cortex predicted slower RTs in non-gamers. While gamers exhibited elevated FA along the left superior occipital–inferior parietal dorsal stream,^23^ neither this tract nor the part of the dorsal stream between the calcarine–occipital FA connection showed a direct behavioral relationship. In contrast, FC between the left superior occipital gyrus and superior parietal lobule was significantly associated with faster RTs. Moreover, SC-constrained analyses revealed that higher local efficiency in the left superior occipital cortex predicted faster responses, whereas greater node degree was associated with slower performance^25^. Additionally, drawing from our SC-dFC (sender) coupling results, the coupling of the number of tracts with the calcarine sulcus was associated with slower response times and was elevated in non-gamers.

Together, these findings suggest that non-gamers rely more heavily on investing cognitive resources into early visual processing, likely putting more effort into object discrimination and resolution of individual dot trajectories during rapid visuomotor decisions to compensate for reduced visuomotor integration. Since the dorsal stream’s structural integrity was previously shown not to be predictive of performance in prior work within the same dataset^23^, the tracts contributing to slower RTs are more likely to be occurring outside of this core visuomotor pathway.^54-56^ This reinforces the interpretation that while early sensory processing is essential, behavioral efficiency is more tightly linked to visuomotor transformation and motor-readied, goal-driven action selection. Cognitive resources appear to yield greater behavioral returns when invested in converting sensory input into action once a satisfactory perceptual benchmark is reached, rather than in the monotonic refinement of early-stage encoding, which exhibits diminishing returns.

Additional behaviorally relevant structural connections reinforce this interpretation. Faster responses were associated with higher axial diffusivity (AD) between the superior temporal pole and inferior frontal orbital cortex regions critical for integrating high-value sensory information^57^ with goal-directed evaluation^58-60^. In this context, the superior temporal pole likely receives scene-specific contextual input from the parahippocampus, enabling the inferior frontal orbital cortex to transform the learned value of this contextual information into action during visuomotor decisions.

Similarly, greater fractional anisotropy (FA) between the parahippocampus and left precuneus was linked to faster RTs. The parahippocampus plays a key role in recognizing scene-specific spatial relationships^61^, while the precuneus supports visuospatial imagery and internal simulation^62-64^—both directly relevant to anticipating movement trajectories based on relative motion cues.

#### 3.3.4 Structure–Function Coupling

Structure–function coupling metrics provided further insight into behaviorally relevant dynamics measuring how well a region’s anatomical infrastructure aligns with its functional load^45^. Both intensive (e.g., FA, AD, NQA, NRDI) and extensive (e.g., count, ncount2) structural metrics were shown to be behaviorally relevant.

In this study, individuals with stronger NDRI–FC and AD–FC coupling in the right mid-occipital region exhibited faster response times. Given this region’s established role in spatial information processing^65^. This behaviorally relevant finding suggests that stronger coupling may reflect greater alignment between incoming visual load and the region’s capacity to relay spatial information to downstream nodes, thereby facilitating response selection. Prior work from our SC-constrained analyses showed that gamers exhibit greater local efficiency in both SC constrained FC and SC constrained dFC networks involving this same region^25^. This suggests that, at equivalent levels of coupling strength, gamers’ networks are more structurally optimized to disseminate spatial information, enabling faster integration with behaviorally relevant areas. Such an architecture may indirectly support more efficient visuomotor transformations, even when the degree of structure-function alignment is comparable between groups.

The coupling of count with sender dFC in the left frontal medial orbital cortex was greater in gamers and negatively correlated with response time, indicating faster behavioral performance. This aligns with earlier findings showing that faster RTs were also facilitated by greater axial diffusivity between the left superior temporal pole and the left inferior frontal orbital cortex. Both medial and inferior frontal orbital regions are implicated in integrating value-based information with action planning ^59, 60^. In this context, this bolsters previous findings that gamers engage in more feedforward, top-down selection of goal-relevant motor responses based on high-level contextual information such as relative motion cues extracted from the scene, including the motion of target dots relative to distractors^25^. The right superior anterior cingulate results were consistent with prior evidence of greater DAN→SN network switching in gamers and the role of superior ACC as a core node in the salience network, SN ^24^. Additionally, as discussed earlier, increased reliance on early visual processing in non-gamers may explain the higher ncount2 coupling with sender dFC in the calcarine R. This finding supports our claim that non-gamers are overly dependent on early-stage visual input, which likely contributes to downstream bottlenecks during visuomotor transformation and action execution—ultimately slowing visuomotor decision-making response times compared to gamers.

#### 3.4.5 Synthesis of Findings Across Modalities

Across modalities, a consistent pattern emerged that faster response times were associated with stronger connectivity in circuits involved in transforming high-value sensory information into goal-directed action and supporting goal-directed motor execution, *i.e.,* regions implicated in resolving perceptual ambiguity more effectively tracked with faster RT. Central to this process was the superior anterior cingulate cortex (ACC sup), a region traditionally associated with conflict monitoring and the resolution of competing motor plans^66^. More recent work suggests the ACC functions as a key hub for uncertainty-driven cognitive control, particularly in dynamic and feedback-sensitive decision environments^67-70^. These areas likely serve to clarify salience signals and commit to goal-relevant actions once monitoring demands have been satisfied. The superior ACC was previously found to be a significant node involved with the increased DAN-to-SN interaction observed in gamers, which tracked with improved RT ^24^ and within the right ACC, a node by which enhanced SdFC sender coupling was tracked with improved response times across multiple SC measures.

Conversely, increased activity among regions involved in motor correction, stress reactivity, and dopaminergic signaling was positively correlated with response time, indicating slower performance and stronger involvement in non-gamers. This pattern may reflect reduced access to high-value salient cues, as suggested by greater engagement of early visual areas and cerebellar–midbrain–thalamic circuits^51, 71, 72^. These compensatory pathways likely reflect increased reliance on bottom-up processing and corrective feedback, with the anterior cingulate cortex (ACC) potentially recruited to manage elevated uncertainty during response selection.

Examples of regions implicated in resolving ambiguity in these findings include the left medial frontal orbital cortex, an executive region critical for integrating value-based information with motor planning. SdFC sender coupling with streamline count in this region was significantly stronger in gamers (*p* = 0.32, *d* = 0.74) and negatively correlated with response time (*r* = –0.34, *p* = 0.035), consistent with improved performance. Another example includes increased synchrony, measured by functional connectivity, between the red nucleus—a lower midbrain structure canonically linked to limb control—and the pulvinar thalamus, reflecting enhanced bottom-up and top-down coordination. This connection likely contributes to perceptual disambiguation by signaling motor readiness^73-75^, effectively facilitating the “I’m ready to press the button” moment.

This finding parallels prior work in Go/No-Go paradigms, where red nucleus activity modulates motor output based on recent trial history, speeding responses after Go trials and promoting caution after Stop trials. Critically, red nucleus neurons amplify directional signals during successful Stop trials, suggesting a role in reshaping ongoing motor plans when initial responses must be inhibited. This capacity for rapid, context-sensitive motor adjustment reflects a form of feedforward control, in which the system anticipates task demands and dynamically tunes motor output in real time. The functional connection between the red nucleus and pulvinar thalamus is supported by a group-level difference favoring gamers (*p* = 0.0061, *q* = 0.02, *d* = 0.85), and negatively correlated with response time (*r* = –0.41, *p* = 0.009), which suggests that long-term exposure to high-stakes, fast-paced environments fosters a neurocognitive strategy that prioritizes feedforward conflict resolution—facilitating swift action and flexible, real-time motor adjustments under uncertainty.

These findings collectively confirm our second hypothesis that a novel, data-driven approach based on rcPCA can successfully identify behaviorally relevant connections and structure–function couplings. This method provided a more granular and meaningful view of the neuroplastic refinements observed in gamers and laid the groundwork for testing our primary hypothesis.

Overall, gamers demonstrate enhanced top-down cognitive clarity, unobstructed translation of learned value into action, and bottom-up motor readiness when making visuomotor decisions under uncertainty. This configuration reduces the need for prolonged internal conflict resolution between competing motor plans and offers what may be a functionally necessary and sufficient explanation for their accelerated decision-making compared to non-gamers.

This neurocognitive visuomotor decision making profile supports a feedforward, proactive strategy for visuomotor transformation, through which gamers exhibit an optimized neural architecture for fast, adaptive, goal-directed behavior—one that more readily anticipates possible task outcomes and enables flexible, real-time motor corrections. By reducing internal uncertainty more efficiently, this yields a more effective cognitive architecture for action selection under dynamic, time-sensitive conditions.

In the context of our modified moving-dots task, this strategy was expressed as greater reliance on top-down selection of goal-directed responses based on scene-specific contextual cues, such as the motion of target dots relative to distractors. Gamers adapted more effectively to either possible outcome of a 50/50 visuomotor decision, reflecting enhanced cognitive flexibility and attentional control.

Our results support the primary hypothesis that long-term action video games (AVG) play reflects neuroplastic refinements that reduce visuomotor surprise by shifting toward superior decision-making strategies—ones that more effectively resolve prediction error and minimize internal conflict regarding competing motor plans. This facilitates a more optimal cognitive state for visuomotor decisions, one that is primed to rapidly incorporate salient, task-relevant information and execute swift, accurate, and decisive actions.

Ultimately, these neuroplastic refinements reflect a reallocation of cognitive resources toward circuits that minimize visuomotor surprise more efficiently. Cognitive Resource Reallocation (CRR) thus emerges once again as a viable mechanistic explanation hypothesized to drive the enhancement of a baseline cognitive state over time, enabling refinements in neural configuration adapted to repeated task demands. This shift toward optimizing internal conflict resolution may not merely reflect a gaming-related adaptation but a broader principle regarding how cognitive systems respond to prolonged cycles of strenuous task engagement and recovery—gradually reallocating resources to stabilize changes in cognitive action over time, promoting local refinements that enhance overall efficiency during task performance by targeting regions and connections most involved in reducing task-induced strain.

In the case of AVG experience, this would lead to neuroplastic refinements that translate into more efficient visuomotor decision-making—a common and frequent demand during gameplay, where errors often come with significant costs. Over time, the brain learns to prioritize high-value visual cues, promote goal-directed action, and increase motor readiness to respond to uncertainty in dynamically changing environments. This ultimately would result in quantifiable improvements, establishing a new set point by which the brain makes visuomotor decisions, leading to the ∼190 ms response time advantage observed in gamers, without any loss in accuracy.

As gamers repeatedly engage in high-stakes, rapid-response tasks, the CRR framework posits that the brain reallocates resources toward pathways that facilitate more proficient AVG performance. These reallocations map cleanly onto cognitive improvements consistently reported in the literature, including enhanced visual acuity^32^, visuomotor integration^11, 23^, attentional control^8^, and cognitive flexibility^12^. The point to be emphasized here is that, rather than endlessly refining early sensory representations beyond a sufficiently reliable threshold, long-term AVG experience encourages the brain to avoid diminishing returns—its priority is to execute the task as efficiently as possible while incurring minimal strain. Over repeated cycles of task engagement and recovery, resources are reallocated toward downstream processes more directly involved in effective gameplay, particularly those supporting visuomotor decision-making and perception–action coupling. This shift offers a comprehensive and mechanistically grounded account of the functional gains observed in gamers and provides a clear neural signature of long-term adaptive plasticity associated with AVG play.

#### 3.4.6 Methodological Strengths, Limitations, and Future Directions

PCA is fully data-driven and agnostic to anatomical priors, enabling it to reveal latent patterns of intersubject variability that might be missed in anatomically constrained pipelines. The rcPCA ROI selection method presented here offers several practical strengths. By identifying relevant ROIs *a priori* based on structured variance, the approach reduces the multiple comparisons burden inherent in full-brain ROI-wise testing. Rather than applying statistical correction across tens of thousands of univariate tests, corrections are reserved for only the most informative regions and their corresponding connections. This not only enhances statistical power but also improves interpretability and replicability, particularly in studies with modest sample sizes.

Our study recruited a sample of healthy young adults, allowing us to isolate the effects of long-term video game playing while minimizing confounds. However, this design choice also imposes certain limitations. First, the dataset in this study captures a cross-sectional snapshot of individuals with long-term action video game experience. As such, it does not permit inferences about the rate of neuroplastic adaptation over time, limiting the ability to make direct causal claims. Determining how these changes evolve would require longitudinal studies and clinical training interventions. Additionally, our gender distribution was not balanced between gamers and non-gamers, limiting our ability to analyze gender-specific differences in brain and behavioral responses. While participants were recruited from university campuses with presumably similar educational backgrounds, we did not explicitly screen for education levels or cognitive ability, meaning we cannot establish direct correlations between baseline cognitive performance and task outcomes.

There, a larger sample size would substantially benefit future research by enabling more sensitive testing of within-group brain–behavior correlations, making it possible to determine whether observed effects are driven primarily by gamers, non-gamers, or both. These distinctions could clarify group-specific visuomotor strategies and reveal potential subtypes of neural adaptation associated with different gaming subgenres. Additionally, larger cohorts would support a more granular analysis of individual differences and enable more robust comparisons across gaming subgenres.^6^

Another limitation is the absence of resting-state fMRI (rs-fMRI) data. Given the success of rcPCA in identifying behaviorally relevant ROIs, it would have been valuable to map these regions onto canonical resting-state networks and compare their connectivity profiles with rs-fMRI data.^76^ This could offer additional insight into how task-based network dynamics relate to intrinsic functional organization. Unfortunately, resting-state scans were not collected as part of this study, limiting our ability to explore these relationships.

A central assumption of rcPCA’s ROI filtering method used in this study was that the dominant axes of intersubject variance, as identified by PCA, reflect functionally or behaviorally meaningful structure. While this assumption held strongly in the present study, yielding reproducible group differences and behaviorally relevant effects across modalities, it may not generalize to all datasets or neuroimaging metrics. PCA’s linear decomposition may fail to fully capture complex variance structures in metrics characterized by nonlinear relationships.

Our empirical validation supports this view. Spearman rank significance was lost when contributions were limited to the top 20 ROIs per component in dFC receiver and SC connectivity metrics but fully recovered when cumulative contributions across all ROIs were used. This suggests that meaningful contributors may be more diffuse and distributed non-monotonically across components. Such discrepancies may indicate more complex or non-linear variance patterns. Future work could apply nonlinear methods like kernel PCA, manifold learning, or tensor decompositions to explore whether residual variance reflects interactions or features that escape linear separation. At a minimum, these results emphasize the value of combining linear methods like PCA with downstream validation to ensure comprehensive coverage of relevant variance space.

ICA may offer further insights in such cases. While ICA is linear, it leverages non-Gaussianity to uncover statistically independent sources of variance, revealing complementary patterns that PCA’s variance-maximizing axes may not capture. Moreover, PCA-derived ROIs could serve as informative priors for joint ICA or other multimodal data fusion frameworks, extending this method’s broader applicability.

Additionally, applying dynamic PCA or tensor decomposition could allow the method to track time-varying or cross-modal shifts in variance structure, offering potential applications in areas such as adaptive brain–computer interfaces (BCIs). Finally, PCA-derived metrics may enhance classification pipelines, supporting both subject-level fingerprinting and group-level identification of neurobiological phenotypes.

Another useful extension would involve calculating the directional skew of each ROI by taking the difference between sender and receiver contributions in dFC (i.e., sender minus receiver). To enable comparisons across subjects and modalities, this difference could be normalized by the total contribution (i.e., sender plus receiver), yielding a normalized score in the range [–1, 1].

Thus, a Directional Skew Index (DI) to capture asymmetries in directed connectivity (dFC) would be defined as

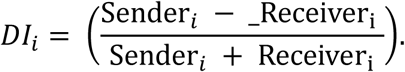

A value near +1 would indicate a strong net sender (information source), –1 a strong net receiver (information sink), and 0 a balanced node with symmetrical inflow and outflow. This normalized skew metric could offer deeper insight into causal asymmetries and dynamic role shifts in directed connectivity. By recovering the directional specificity otherwise collapsed in total influence measures, it enhances interpretability and enables a more nuanced characterization of regional dominance in information flow, particularly in the absence of strong priors.

This method offers ample room for future development. Incorporating demographic or clinical covariates (e.g., age, sex, IQ, education, symptom scores) and genetic markers such as single-nucleotide polymorphisms (SNPs) could help contextualize sources of variance and improve generalizability across populations. With further refinement, these tools could contribute to the development of diagnostic or prognostic frameworks in clinical neuroscience. Finally, benchmarking the PCA-derived ROI contributions against alternative feature selection strategies, including ICA, clustering algorithms, or model-derived importance scores from machine learning pipelines, may further clarify when and where this approach provides the most utility. Evaluating robustness across large, multi-site datasets is also critical for assessing generalizability and translational potential. Taken together, the PCA-based ROI selection framework offers a scalable, mathematically principled approach to dimensionality reduction of brain regions to those most informative as measured by contribution to explained variance in neuroimaging analysis. Expanding the framework to include pairwise ROI coupling or grouping ROIs into resting-state or task-defined networks may enable richer interpretations at the level of functional systems rather than individual regions

While rooted in a linear decomposition, it opens the door to nonlinear and multimodal extensions and serves as a flexible foundation for developing future tools that can adapt to the complexity and diversity of modern brain data.

#### 3.4.7 Conclusion

This study introduces a novel, data-driven PCA-based method for ROI selection that reveals significant neuroplastic adaptations plausibly driven by action video game (AVG) experience. Our findings provide empirical support for the Cognitive Resource Reallocation (CRR) framework, which proposes that the brain optimizes performance by reallocating functional and metabolic resources toward anatomically plausible, behaviorally relevant circuits to more effectively meet arduous task demands—ultimately striving to reduce costly prediction errors. Our findings show that AVG experience may promote more efficient visuomotor decision-making through top-down cognitive clarity, the unobstructed transformation of learned value into goal-directed action, and bottom-up motor readiness. The convergence of these factors would effectively reduce internal conflict, mitigate visuomotor surprise, and enable rapid yet skillful action selection through greater anticipation of multiple outcomes in the face of uncertainty.

These findings deepen our understanding of the neural mechanisms supporting enhanced visuomotor performance in gamers and have broader implications to inform educational strategy, improve learning programs, and better facilitate skill acquisition. The ability to more effectively reduce uncertainty and resolve internal conflict in high-pressure contexts—as observed in gamers—could potentially inform future design of educational programs, rehabilitation protocols, and high-stakes professional training, including in sports, surgery, and military operations. Moreover, these findings may help shape macro-level rehabilitation goals, especially for patients relearning motor coordination and decision-making in dynamic environments.

Our rcPCA method offers ample room for growth and future expansion, such as integrating demographic or clinical variables or applying nonlinear approaches like kernel PCA to capture more complex variance patterns. This work not only demonstrates the utility of data-driven, principled neuroimaging methods but also provides a foundation for research aimed at enhancing human performance through targeted adaptation.

While future work should assess the generalizability and nonlinear potential of rcPCA, the present study offers both a novel analytical tool and meaningful insight into how experience reshapes cognition. It highlights the brain’s remarkable ability to "level up" by repeatedly confronting and striving to overcome cognitively demanding challenges. Through CRR, these challenges become less effortful over time, minimizing conflict, reducing uncertainty, and ultimately enabling more efficient, high-performance behavior. This study not only confirms that action video game play reflects targeted neuroplasticity but also offers a generalizable blueprint for cognitive optimization through adaptive resource reallocation.

## 4. Materials and Methods

### 4.1. Subject Data

A total of 47 right-handed participants were recruited for this study, including 28 action video game players (gamers; 4 female) and 19 non-gamers (12 female). The groups were age-matched (gamers: 20.6 ± 2.4 years; non-gamers: 19.9 ± 2.6 years). Participants were classified as "gamers" if they reported playing at least five hours per week of one or more action video game genres—including First-Person Shooter (FPS), Real-Time Strategy (RTS), Multiplayer Online Battle Arena (MOBA), or Battle Royale (BR)—consistently over the past two years. Non-gamers reported playing less than 30 minutes of video games per week over the same period. All participants passed the Ishihara color vision test to confirm normal color perception. A modified left–right moving-dot (MD) task was used to probe visuomotor decision-making and assess group differences in response time and accuracy.

After preprocessing and quality control, the final sample size varied slightly by modality. Functional and directed functional connectivity (FC and dFC) analyses included 43 participants (25 gamers and 18 non-gamers), while brain–behavior correlations using FC and dFC included 41 participants (24 gamers and 17 non-gamers). Structural connectivity (SC) analyses included 46 participants (27 gamers and 19 non-gamers), with 44 participants (26 gamers and 18 non-gamers) used in SC brain–behavior analyses. Finally, SC–FC and SC–dFC coupling analyses included 42 participants (24 gamers and 18 non-gamers), and the corresponding brain–behavior coupling analyses included 40 participants (23 gamers and 17 non-gamers).To confirm eligibility and assign participants to the appropriate group, a questionnaire was administered assessing video game genre and play frequency over the past two years. All participants passed the Ishihara Test for Color Deficiency and completed informed consent and health screening forms before data collection. The study was approved by the Institutional Review Boards of Georgia State University and the Georgia Institute of Technology, both located in Atlanta, Georgia.

### 4.2. MRI Data

#### 4.2.1 Data Collection, Scanning & Tractography Protocols

Whole-brain structural and functional MR imaging was conducted on a 3T Siemens Magnetom Prisma MRI scanner (Siemens, Atlanta, GA, USA) at the joint Georgia State University and Georgia Institute of Technology Center for Advanced Brain Imaging, Atlanta, GA, USA. High-resolution anatomical images were acquired using a T1-MEMPRAGE scan sequence for voxel-based morphometry and anatomical reference. The acquisition parameters were as follows: TR = 2530 ms, TE1-4 = 1.69–7.27 ms, TI = 1260 ms, flip angle = 7°, and voxel size = 1 mm × 1 mm × 1 mm.

Diffusion-weighted imaging (DWI) data were collected using a multi-shell diffusion scheme with b-values of 300, 650, 1000, and 2000 s/mm², corresponding to 4, 17, 39, and 68 diffusion-encoding directions, respectively. One non-diffusion-weighted (b = 0) volume was also included. The acquisition was performed using a single-shot echo-planar imaging (EPI) sequence with anterior-to-posterior (AP) phase encoding. Each diffusion volume consisted of 60 axial slices acquired with a 2 mm isotropic resolution (slice thickness = 2 mm, in-plane resolution = 2 × 2 mm), and the field of view (ReadoutFOV) was 220 mm. The total scan duration was approximately 6.5 minutes. Acquisition parameters for the diffusion imaging included TR = 2750 ms and TE = 79 ms.

Following data acquisition, the diffusion data were reconstructed in the MNI space using q-space diffeomorphic reconstruction (QSDR)^77^ to obtain the spin distribution function (SDF)^78^ in DSI Studio (Version Hou, 2024). A diffusion sampling length ratio of 1.25 was applied, with the output resolution in diffeomorphic reconstruction set to 2 mm isotropic. The tensor metrics were then calculated.

For fiber tractography, a deterministic fiber tracking algorithm^79^ was used, incorporating augmented tracking strategies^80^ to improve reproducibility. The quantitative anisotropy (QA) threshold was set to 0.12, and the angular threshold was set to 60 degrees. The step size was 1.00 mm, and tracks shorter than 10 mm or longer than 400 mm were discarded. A total of 5 million tracts were calculated for each subject. Shape analysis was conducted to derive shape metrics for the tractography^80^

For full reproducibility, the parameter ID used in DSI Studio to configure these settings is 8FC2F53D9A99193Fba3Fb803Fcb2041bC843404B4Cca01cbaCDCC4C3Ec. This ID allows others to load the exact settings and parameters used in our analysis, ensuring that the tractography and other metrics can be reproduced using the same configurations.

The functional imaging was performed using a T2-weighted gradient echo-planar imaging (EPI)* sequence during the behavioral tasks. Four functional runs were acquired with the following parameters: TR = 535 ms, TE = 30 ms, flip angle = 46°, and voxel size = 3.8 mm × 3.8 mm × 4 mm. The field of view was 240 mm, and 32 slices were collected in an interleaved order with a slice thickness of 4 mm. A total of 3440 brain images were acquired during task performance.

#### 4.2.2 fMRI Pre-Processing Pipeline

The preprocessing pipeline for functional MRI data combined with tools from AFNI ^81, 82^ and FSL^83-85^ to ensure high-quality data for subsequent analysis.^86-88^ The process began with denoising the fMRI data to reduce noise from physiological artifacts such as head motion and scanner drift, using AFNI’s dwidenoise command. This step resulted in denoised datasets for each run of the fMRI data. Following this, motion correction was performed using AFNI’s 3dvolreg, which registers each volume of the fMRI data to a reference volume within the session. After motion correction, the data were aligned to the MNI space using FSL’s FLIRT tool, ensuring that all data were in a common standard space for group-level comparisons.

To remove any motion-related artifacts, outlier detection was carried out using AFNI’s 3dToutcount, which computes the fraction of outlier voxels in each volume. A censoring procedure was then applied, excluding volumes where the fraction of outlier voxels exceeded a predefined threshold (0.1). Despiking was performed with AFNI’s 3dDespike, which removed brief, spurious signal fluctuations, or "spikes," from the data. Following despiking, slice timing correction was applied using 3dTshift to ensure temporal alignment across slices in each volume.

Time series were then extracted from predefined brain regions based on the AAL3 parcellation using AFNI’s 3dROIstats, which computes the average signal within each region of interest (ROI). These time series were saved as text files for further analysis. Signal-to-noise ratio (SNR) for each run was computed using AFNI’s 3dTstat to calculate the mean signal and 3dTproject to compute the standard deviation of the noise. Additionally, global correlation averages (GCOR) were computed to assess overall data quality.

The degree of spatial blurring in the data was estimated using FSL’s 3dFWHMx, which calculates the full width at half maximum (FWHM) of the data’s spatial blurring. Following this, an extents mask was created using AFNI’s 3dmask_tool to identify valid brain regions with usable data across all volumes. The data were then registered to the MNI template (ENIGMA Template) using AFNI’s @auto_tlrc, which applies a transformation matrix to warp each subject’s data into standard space for group-level analysis.

To further improve data quality, Principal Component Analysis (PCA) was applied using AFNI’s 3dpc to remove non-neuronal signals, such as global signal fluctuations and motion-related noise. PCA regressors were generated from ventricular and brain regions and were used in subsequent regression analysis to remove unwanted variance from the data. Finally, the processed datasets were reviewed for quality control, and any remaining temporary files were removed to prepare the data for further analysis.

After the AFNI and FSL preprocessing steps, additional processing was performed in MATLAB to further refine the data for task-based analysis. This included outlier correction, where extreme values in the time series that exceeded 5 standard deviations were identified and corrected. Detrending was applied to remove any linear trends from the data using MATLAB’s detrend function, ensuring that any slow drifts in the signal did not affect subsequent analyses. The time series data were then parsed by behavioral condition and time block, creating condition-specific time series data for each subject. This allowed for a more detailed analysis of brain activity that aligned with specific experimental conditions. The preprocessed time series data for each region of interest (ROI) in the AAL3 atlas were stored in structured files and saved for subsequent analysis.

### 4.3 Atlas Selection & AAL3 Parcellation

For the functional and structural connectivity analysis, the Automated Anatomical Labeling 3 (AAL3) atlas was selected due to its widespread use and strong effect sizes in capturing brain structure-function relationships, especially when compared to other well-known atlases.^89^ The AAL3 atlas includes 166 parcellations, with critical task-relevant regions such as the orbitofrontal cortex, cerebellum, and thalamic nuclei, which are particularly relevant for video game studies investigating neural processes underlying cognitive functions like visuomotor decision-making. This makes it highly suitable for whole-brain analysis in our study.^38^ For better visual clarity and interpretability, we organized the regions of the AAL3 atlas into clear subdivisions Orbitofrontal, Occipital, Limbic System, Frontal, Temporal, Thalamus, Parietal, Basal Ganglia, Cerebellum, and Brain Stem (Figure 7, a) based on their known anatomical locations, while preserving individual regions in our analysis as displayed in Supplementary Figure 1. As the brain slices progress in (Figure 7, b), the organization of these regions becomes more apparent, revealing how these anatomical structures are spatially arranged. This clear organizational structure aids in interpreting the results of our analysis.

**Figure 7.**
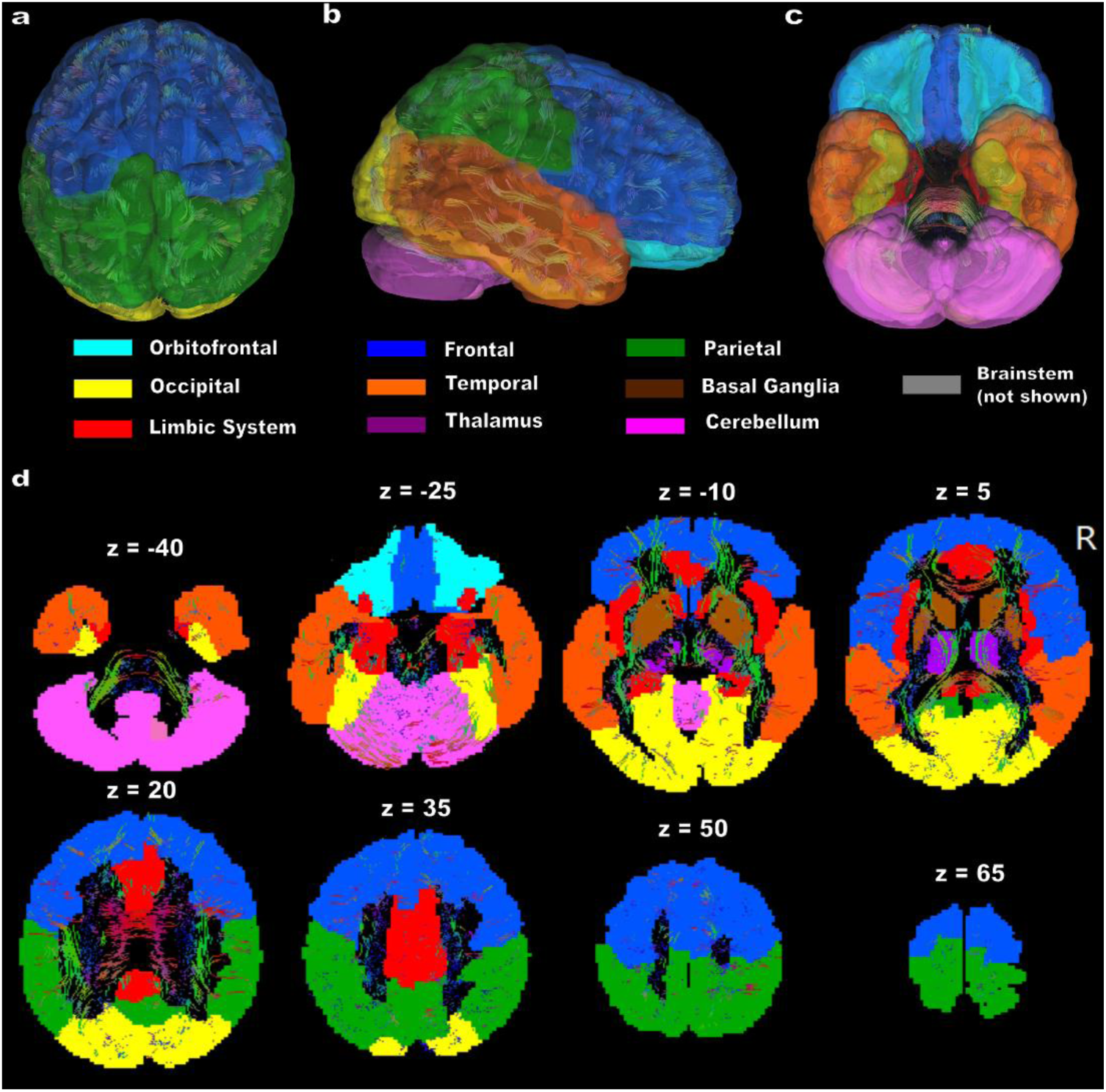
AAL3 Atlas Parcellation Categories for Connectivity Analysis. Visualization of the AAL3 atlas with anatomically grouped parcellations used in the connectivity analysis. **(a)** Superior view, **(b)** Right lateral view, **(c)** Inferior view. **(d)** Axial slices illustrate the parcellation structure along the z-axis. Colors correspond to distinct anatomical groups. The brainstem (gray) is not shown but is included in the analysis.

### 4.4 rcPCA-Based ROI Selection for Connectivity and Structure-Function Coupling Analysis

#### 4.4.1 Connectivity and Coupling Data

After preprocessing, functional connectivity (FC) was computed using pairwise Pearson correlations across the full set of parcellated time series (AAL3 regions). Directed functional connectivity (dFC) was estimated using pairwise time-domain Granger causality, following the procedure outlined in Dhamala et al.^90^ The evaluation of TGC was conducted in the frequency band in the range between *f*_1_= 0.05 Hz to *f*_2_ = 0.9 Hz, with a sampling rate of 1.87 Hz (*TR*^−1^).

The appropriate model order for the TGC analysis was determined by minimizing the spectral difference between the Granger-generated time series and the original signal, while maintaining sensitivity to trial-specific dynamics governed by the trial duration and the repetition time (TR). To preserve this sensitivity, the model order was constrained such that it did not exceed the number of time points within a trial. The maximum allowable model order, denoted mo_max_, is given by

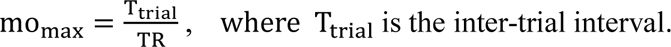

The model order of 5 was selected as it best minimized the spectral difference in our dataset while preserving the sensitivity required to accommodate trial-specific GC influences. GC matrices were computed between the time series of the AAL3 brain regions, using this optimal model order, which effectively became our directed functional connectivity measure for this study.

For structural connectivity (SC) analysis, diffusion-weighted imaging data were processed to derive a structural connectivity matrix based on deterministic tractography using DSI-Studio. This process involved the use of q-space diffeomorphic reconstruction (QSDR)^78, 80^ and a deterministic fiber tracking algorithm^79^ to estimate the structural connectivity between brain regions. The resulting matrix quantified the number and strength of the structural connections between each pair of brain regions. To assess the degree of alignment between anatomical structure and functional signaling, structure–function coupling (SFC) metrics were computed by correlating structural connectivity (SC) matrices with both undirected functional connectivity (FC) and directed functional connectivity (dFC).^45^ All matrices were aligned using the AAL3 parcellation to ensure consistency across modalities.

The coupling between SC and FC (SFC) was computed by extracting, for each ROI pair, the corresponding values from the SC and FC matrices. Pearson correlation was then used to quantify the strength of association between anatomical connectivity and functional co-activation across subjects. The same approach was used to compute coupling between SC and directed functional connectivity (SdFC), where each subject’s SC matrix was compared against sender-mode Granger causality matrices. Although SC is symmetric and undirected, the correlation was performed by holding the row index constant and correlating each region’s structural projection pattern with its outgoing directed functional influences. This resulted in a region-wise measure of SdFC coupling, reflecting how well the structural architecture of a region supports its role as a functional sender.

Both intra-regional and pairwise coupling values were computed; however, to maintain a focused and interpretable analysis in the current study, we limited our exploration of structure–function coupling to intra-regional sender-mode values. This approach allowed us to establish a clean, region-specific profile of structure–function alignment. Future work may extend this framework to include full pairwise coupling analyses as a means of capturing distributed patterns of structural–functional integration.

#### 4.4.2 rcPCA-Based ROI Selection

To identify the most informative regions of interest (ROIs) across structural and functional connectivity domains, we developed a data-driven, region-cumulative principal component analysis (rcPCA) method for dimensionality reduction. The rcPCA approach decomposes subject-level connectivity matrices into orthogonal principal components and derives regional contribution scores by quantifying each ROI’s influence on the structured variance of the dataset. All analyses were implemented in MATLAB using custom scripts developed for this study. This method adapts standard PCA protocols and interpretation techniques commonly used in the neuroimaging community, focusing on high-loading features per component to enable principled ROI selection and structured dimensionality reduction without sacrificing interpretability. To prioritize meaningful contributors while minimizing noise, we capped contribution accumulation at the top 20 ROIs per component. This strategy was chosen to avoid rank dilution from low-weight contributors, improve interpretability, and emphasize ROIs that consistently explain variance across components. Including all ROIs per component would risk amplifying weak or spurious contributions, particularly in sparser or asymmetric modalities such as structural or directed connectivity. While this approach may sacrifice global monotonicity with raw variance rankings, it enhances the salience of high-impact regions, aligning with our goal of identifying interpretable, behaviorally relevant ROIs.

For undirected connectivity data such as functional connectivity (FC), we first removed diagonal elements from each subject’s 166 × 166 ROI-wise matrix and reshaped the resulting 3D array into a two-dimensional matrix of size n × E, where n is the number of subjects and E is the number of unique off-diagonal connections between ROIs. This matrix reshaping was handled by the script undirected_pca_analysis.m. This script then called our custom function run_pca.m that uses MATLAB’s built-in *pca()* function, which performs singular value decomposition (SVD)^46^ on a mean-centered data matrix of size n × E. This function returns the component loadings, subject scores, and variance explained by each principal component. ROI-level contributions were used to interpret latent components, while subject-level scores served as PCA-derived features for potential classifier enrichment.

Finally, we identified the ROI contributions for each component by summing the absolute values of the PCA loadings associated with each region. For connectivity-wise decompositions (e.g., FC), loadings were reshaped into full ROI × ROI matrices with NaNs along the diagonal to exclude self-connections. The sum of absolute values across each row yielded a scalar contribution score per region, reflecting how strongly a given ROI influenced the variance captured by that component. For ROI-wise decompositions (e.g., structure–function coupling), each coefficient directly corresponded to a region, and the absolute values were taken as contribution scores. ROIs were ranked in descending order for each component and saved for downstream analysis.

To compute each region’s cumulative contribution to the explained variance, we used the script *undirected_pca_roi_contributions.m*. This script sets the number of top pca rois, cumulative variance % and calls the custom function *pca*_*roi_contributions.m* which calculates calculates the cumulative contirbution of each roi to the user’s desired explained variance threshold which we set to 80% using a weighted sum set by the weights of each PC By capturing 80% of the total variance across components, we retain the dominant patterns in the data that are most likely to reflect structured signal rather than noise, while discarding components that account for only a small portion of the variance which may be less interpretable and more sensitive to measurement error. This threshold also stabilizes the ranking of contributing ROIs across modalities and ensures that our contribution scores are derived from a meaningful, high-variance subspace rather than dominated by noisy low-variance dimensions. This procedure was carried out not only for FC, but also for all 13 categorically distinct structural connectivity (SC) metrics—(axial diffusivity (AD); streamline count; fractional anisotropy (FA); isotropy (ISO); mean diffusivity (MD); mean streamline length; Ncount; Ncount2; normalized quantitative anisotropy (NQA); non-restricted diffusivity imaging (NRDI); radial diffusivity (RD); restricted diffusivity imaging (RDI))—as well as for all 13 SFC and all 13 SdFC coupling matrices, using the same run_pca.m and ROI contribution framework. For definitions of each metric, see Table 1. ^91^

**Table 1.** Structural connectivity (SC) measures and definitions reflecting standard interpretations of diffusion and tractography measures from DSI Studio.

| SC Measure | Description |
| --- | --- |
| Axial Diffusivity (AD) | Diffusion along the primary fiber axis; reflects axonal integrity. |
| Count | Raw number of tractography streamlines between regions. |
| Fractional Anisotropy (FA) | Degree of directional water diffusion; higher values indicate organized fiber structure, broadly linked to myelination. |
| Isotropy (ISO) | Uniformity of diffusion in all directions |
| Mean Diffusivity (MD) | Average diffusion in all directions; reflects overall water mobility. |
| Mean Length | Average length of streamlines between ROIs |
| None-Restricted Diffusion Imaging (NRDI) | Measures extracellular diffusion; sensitive to edema or extracellular space. |
| Normalized Count (Ncount) | Streamline count adjusted for ROI size and distance. |
| Normalized Count (Inverse-Length Weighted) (Ncount2) | Streamline count weighted by inverse length; emphasizes shorter, potentially more reliable tracts. |
| Normalized Quantitative Anisotropy (NQA) | Tracks the anisotropy of the principal fiber direction, normalized to background noise. |
| Quantitative Anisotropy) QA | Signal intensity along the principal fiber orientation; related to tract integrity. |
| Radial Diffusivity (RD) | Diffusion perpendicular to the main axis; associated with myelin integrity. |
| Restricted Diffusion Imaging (RDI) | Estimates intracellular diffusion; linked to axonal density |

Each analysis followed the same pipeline: data reshaping, PCA decomposition, ROI-wise contribution scoring, and cumulative variance weighting, providing a unified approach to data-driven region selection across modalities.

For directed connectivity data, such as Granger-causal (dFC) matrices, we developed a parallel pipeline using the scripts *directed_pca_analysis.m* and *run_dpca.m*. Since directed matrices represent asymmetric interactions between regions (i.e., sender to receiver), we performed PCA separately for sender, receiver, and total connectivity modes. Directed connectivity matrices are square, where each element represents the strength of influence from one region (sender) to another (receiver). A companion script to *pca_roi_contributions.m*, titled *directed_roi_contributions_dpca.m*, was created to compute cumulative contributions in sender, receiver, and total modes. For each region, cumulative sender influence was calculated by holding the row index constant and summing across columns, whereas cumulative receiver influence was computed by holding the column index constant and summing across rows. Total contributions were obtained by summing both sender and receiver values for each region. These direction-specific contributions were then used to rank ROIs according to their influence in the structured variance of the directed connectivity data.

All outputs—including PCA loadings, explained variance, region rankings, and component-wise visualizations— were saved for each subject group and connectivity modality. In addition, the top ROIs, their corresponding AAL3 region labels, and all cumulative ROI contribution scores were saved to support downstream interpretation and reproducibility. After confirming the internal validity of our approach by comparing PCA-derived region rankings to raw variance rankings within each connectivity metric, these results were used in downstream analyses to identify high-variance ROIs for group comparisons and behavioral correlation testing.

Overall, this rcPCA framework offers a principled and scalable strategy for prioritizing informative brain regions in high-dimensional neuroimaging data. By reducing noise and redundancy while preserving structured variability, the method improves sensitivity to behaviorally relevant effects and significantly lowers the burden of multiple comparisons.

#### 4.4.3 Validation of rcPCA-Derived Region Rankings

To evaluate the internal consistency and robustness of rcPCA, a multi-pronged validation was developed to test the derived regional rankings compared to the raw variance using a combination of rank correlation, permutation testing, and hypergeometric overlap statistics using a significance threshold of *p* < 0.001. This procedure was implemented in the script *validate_pca_variance.m* and carried out separately for each modality, including FC, dFC, SC, SFC, and SdFC(sender).

First, the regions were ranked according to their raw pre-PCA variance. This ranking was then compared to PCA-derived rankings using Spearman’s rank correlation across a range of top-k values, from 2 up to 166, reflecting the full resolution of the AAL3 parcellation. These tests provided converging evidence that the region-based cumulative PCA (rcPCA) method reliably identifies regions based on structured variance patterns, supporting the method’s selectivity and stability across modalities. The choice of k = 20 offered a practical compromise, large enough to capture meaningful connectivity patterns while remaining selective enough to highlight informative ROIs.

In the Spearman analysis, we calculated rank correlation coefficients between the top-k PCA-derived ROIs and the top-k raw-variance ROIs at each k. This approach was chosen for its robustness to non-normal distributions and its sensitivity to monotonic relationships. The correlation coefficient was calculated as

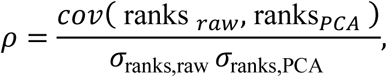

where ranks _raw_ are the ranks of the raw data, *r*anks _PCA_ are the ranks of regions based on the cumulative PCA-derived variance contributions, *cov* is the covariance between two ranked vectors and *σ* is the standard deviation of ranks. The correlation was computed in MATLAB using the *corr()* function with the ‘Type’, ‘Spearman’ option, which is given by

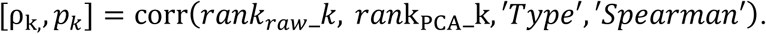

To assess whether the number of overlapping ROIs between the top-k PCA-selected and raw variance-selected regions could be attributed to chance, we used the cumulative distribution function of the hypergeometric distribution. This distribution models the probability of x or more successes (i.e., overlapping ROIs) when drawing two sets of size k from a population of N=166 ROIs without replacement. The probability of observing at least x overlaps under the null hypothesis of random selection is given by the standard expression for hypergeometric p-value

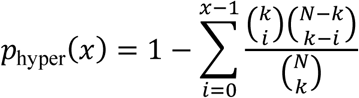

where N is the total number of ROIs (166), k is the number of top-ranked ROIs selected in each set, and x is the observed number of overlapping ROIs between the PCA-based and raw variance-based rankings. This computation was implemented in MATLAB using the built-in hygecdf function and is given by (p_hyper = 1 - hygecdf(x - 1, N, k, k)).

Finally, a null distribution of random overlap values was generated by permuting the raw variance ROI rankings 10,000 times. For each value of k, the top-k ROIs were selected from each permutation, and their overlap with the PCA-selected top-k ROIs, denoted as x, was recorded. The empirical p-value was then computed as the proportion of permutations in which the number of overlapping ROIs was greater than or equal to x, relative to the total number of permutations

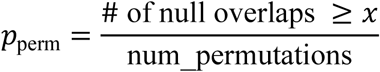

and the equivalent MATLAB statement used for this calculation is given by (perm_pvals(i) = mean(rand_overlaps >= overlap_counts(i))). This allowed us to assess whether the observed overlap between PCA-selected and raw variance-based regions exceeded what would be expected by chance.

All results, including overlap counts, p-values (Spearman, hypergeometric, and permutation), and rank correlations, were logged and plotted across k-values. Summary figures included correlation curves and significance levels as a function of k, bar plots comparing top-k PCA- and raw-variance ROI values, and permutation histograms and hypergeometric threshold overlays for observed overlap. Across all modalities, hypergeometric and permutation tests yielded exceptionally low p-values, often well below the p < 0.001 threshold. At higher values of k, both tests frequently reached the limits of machine precision. In such cases, MATLAB returned literal zero values due to numerical underflow, requiring the imposition of a floor at *p* = 1e-15. This indicated that the observed overlaps were so unlikely under the null distribution that their probabilities could not be accurately represented in double-precision floating-point arithmetic.

This provides compelling evidence that the overlap between PCA-derived and raw-variance ROI rankings was highly unlikely to occur by chance. They further validate the selectivity, robustness, and internal consistency of the proposed method. Notably, using cumulative contributions across all ROIs before selecting the top 20 restored the Spearman rank significance that was lost in the dFC receiver, SC, SFC, and SdFC modalities when only the top 20 ROIs per component were used during accumulation. These validation results confirmed that PCA-selected ROIs consistently overlapped with high-variance regions across modalities, supporting the interpretability, reproducibility, and robustness of the rcPCA selection method. For final visualizations, we used ‘n’ in place of ‘k’ to denote the number of top-ranked ROIs, as it provided a clear and intuitive shorthand in contexts where there was no conflicting N variable.

Furthermore, full convergence of PCA-derived ROI rankings with raw variance rankings, across all validation metrics, was observed within the range of k=100-150. This included Spearman correlations approaching unity, permutation-based p-values below machine-level significance (*p* < 1e-15), and hypergeometric overlaps exceeding chance across all thresholds. Isotropy was the last measure to show full convergence, but did so by k = 90, making the k = 100-150 range a conservative benchmark for achieving maximal rank agreement. This defines a convergence ceiling for PCA-based variance decomposition in whole-brain neuroimaging data for the AAL3 atlas. The observed convergence window likely reflects the resolution of the AAL3 atlas’s 166 ROIs. While this range defines full convergence for this specific parcellation, the maximum k required for machine-level agreement is expected to scale proportionally with atlas dimensionality.

Another observation was that the permutation-based p-values exhibited a characteristic rebound at high k values (e.g., k ≥ 160), as the overlap between random samples and observed sets approached the full ROI space. This behavior reflects the design of permutation tests, which become less discriminative as sampling exhausts the comparison space. This plateau confirms that p-value inflation near the maximum ROI count is an expected property of null model behavior due to sampling saturation.

Together, these results confirm that the PCA-based ROI selection framework reliably identifies regions that meaningfully contribute to structured variance across modalities. The strong convergence across multiple statistical tests affirms that top-ranked ROIs are not artifacts of random variation, but rather reflect well-founded, mathematically principled selection criteria. In short, the method behaved exactly as intended, yielding high-variance-contributing, interpretable ROI candidates for downstream analysis.

#### 4.4.4 Group Comparisons and Brain–Behavior Relationships (rcPCA-Derived ROIs)

To investigate group-level differences and behavioral relevance of connectivity across modalities, we used the top 20 PCA-derived ROIs from each modality-specific decomposition as filters. For each modality—functional connectivity (FC), directed connectivity (dFC; sender, receiver, and total), structural connectivity (SC), and coupling metrics (SFC and SdFC)—we extracted submatrices containing any connection involving a top-ranked ROI. These filtered matrices were used to run non-parametric group comparisons (Mann–Whitney U, rank sum^92^ tests) between gamers and non-gamers and Pearson brain-behavior correlations with response time (RT) as a behavioral measure.

False discovery rate (FDR) correction was applied independently within each connectivity modality to account for multiple comparisons during group comparisons. Two standard approaches were considered: the Storey-Tibshirani (ST) method^93^ (which estimates the proportion of true null hypotheses, π₀) and the Benjamini–Hochberg (BH) procedure^94^ (which ranks p-values and applies a fixed step-up threshold). The choice between them was guided by both observed performance and the underlying assumptions of each method.

The ST method was deemed valid for functional connectivity (FC) data only. In all FC comparisons, the estimated q-values remained consistently less than or equal to the corresponding uncorrected p-values, indicating that the proportion of true nulls could be reliably estimated and that the method behaved as expected. These results suggest that Storey’s assumptions were met for FC—likely due to the dense, symmetric, and continuous nature of FC matrices, along with their relatively uniform distribution of connectivity values.

In contrast, Storey’s method was not considered reliable for the other modalities. In structural connectivity (SC), directed connectivity (dFC), and coupling metrics (SFC, SdFC), q-values were frequently observed to fall below their corresponding p-values despite p-values being well above the uncorrected significance threshold, a pattern inconsistent with valid FDR correction. This behavior indicated instability in π₀ estimation, likely due to skewed or zero-inflated distributions, sparse matrices, and directional asymmetries—conditions known to violate the assumptions behind Storey’s estimator.

To ensure robust and conservative control of false discoveries in these cases, the BH procedure was applied instead. BH does not rely on π₀ estimation and is more stable under non-ideal conditions. It was, therefore, used for all SC, SFC, SdFC, and dFC variants, where structural constraints or directional signal flow introduced potential sources of bias.

For brain-behavior correlations, we computed Pearson correlations between response time (RT) and connectivity values for all connections involving the top 20 PCA-derived ROIs per modality. These correlations were tested with a significance threshold of α = 0.05, and results were further filtered using an effect size threshold of |r| ≥ 0.2 to ensure robustness and interpretability.

---

**SUPPLEMENTARY FIGURE 1.**
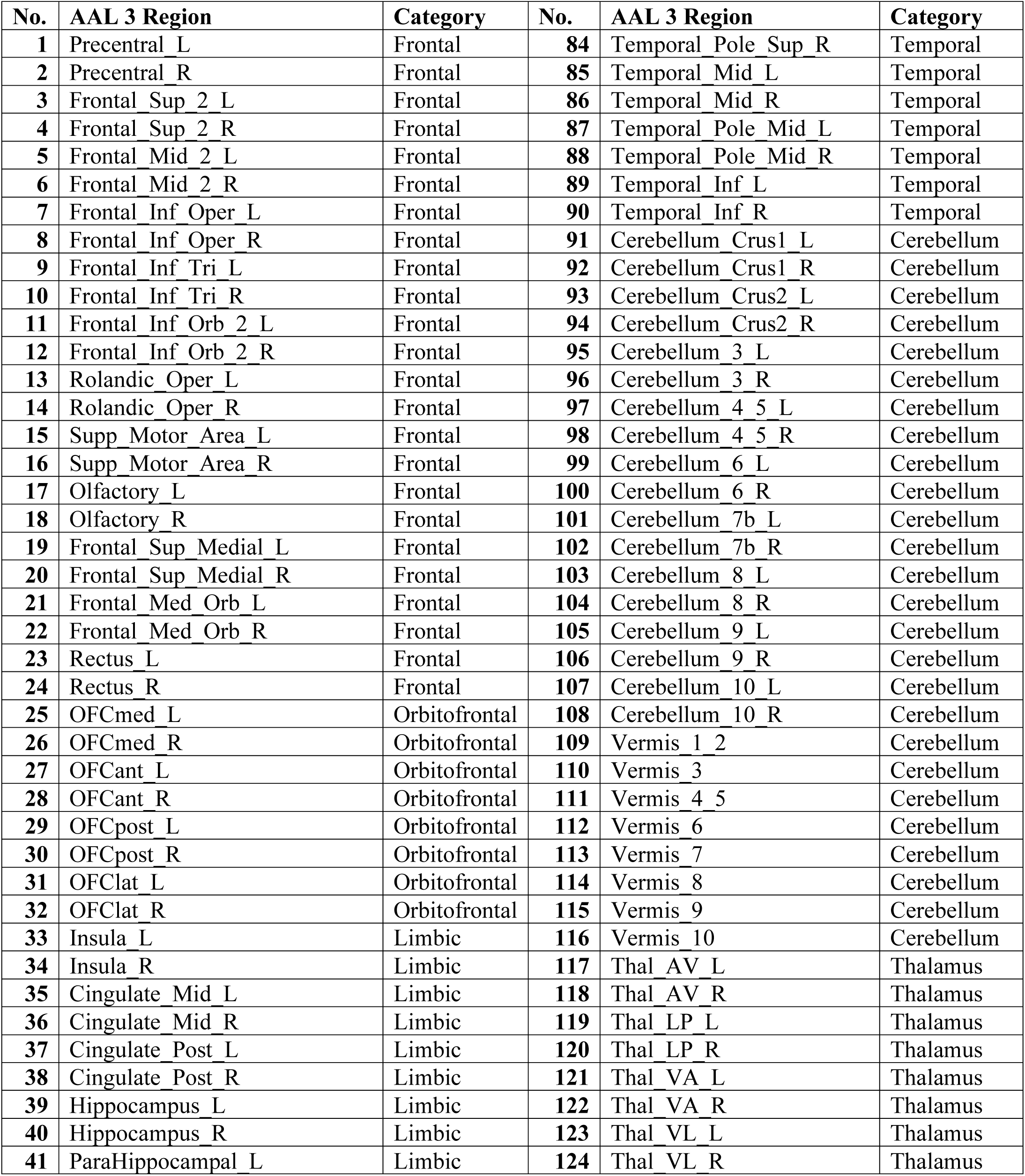
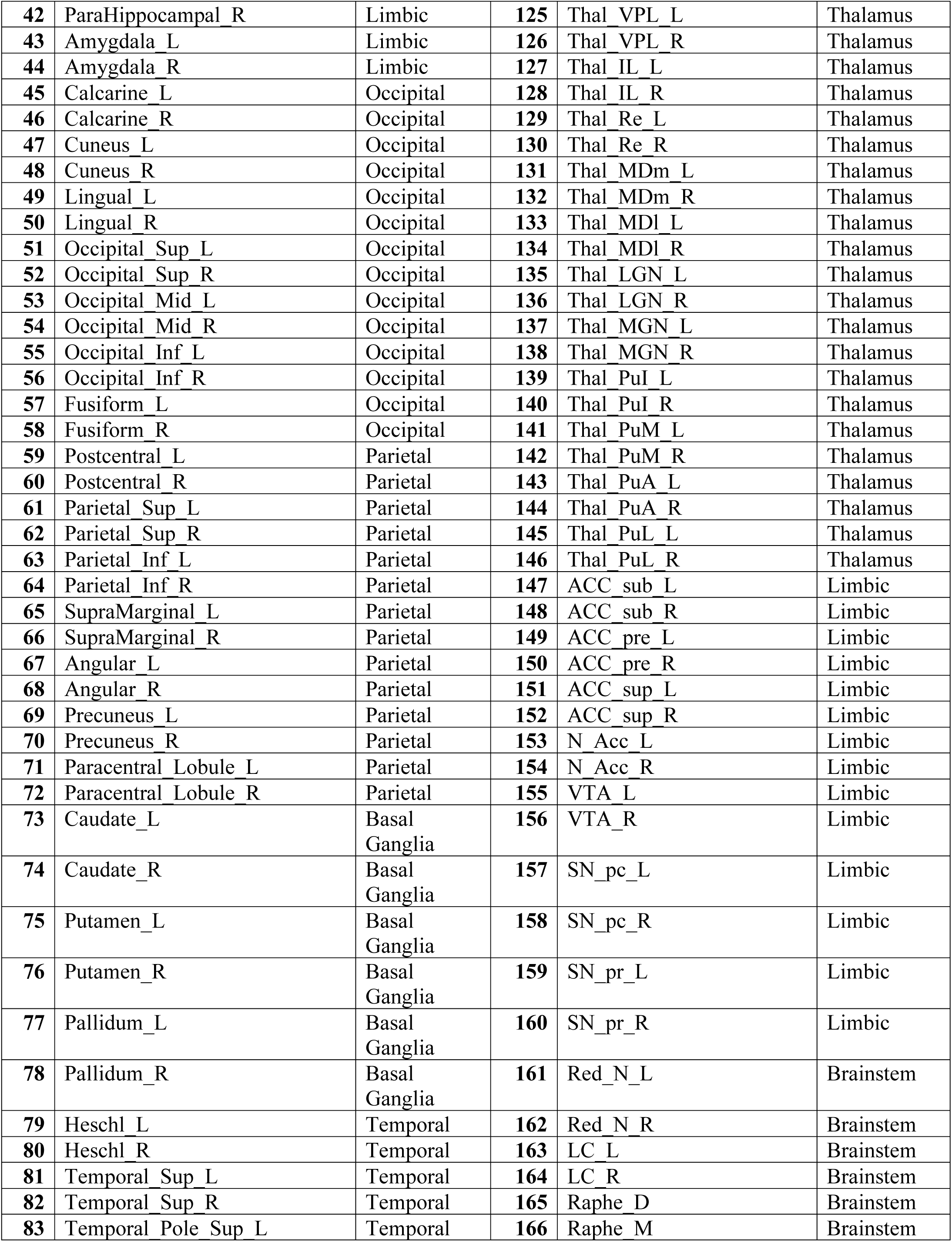
AAL 3 Region Parcellation Categories by Major Anatomical Groups. Original numbers in AAL2 for the anterior cingulate cortex (ACC) and thalamus are left empty in AAL3, as those voxels were substituted by the new subdivisions for the Thalamic nuclei and ACC.^38^ These regions are removed here, and the index has been adjusted accordingly to the correct 166 total parcellations categorized by major anatomical groups.

**SUPPLEMENTARY FIGURE 2.**
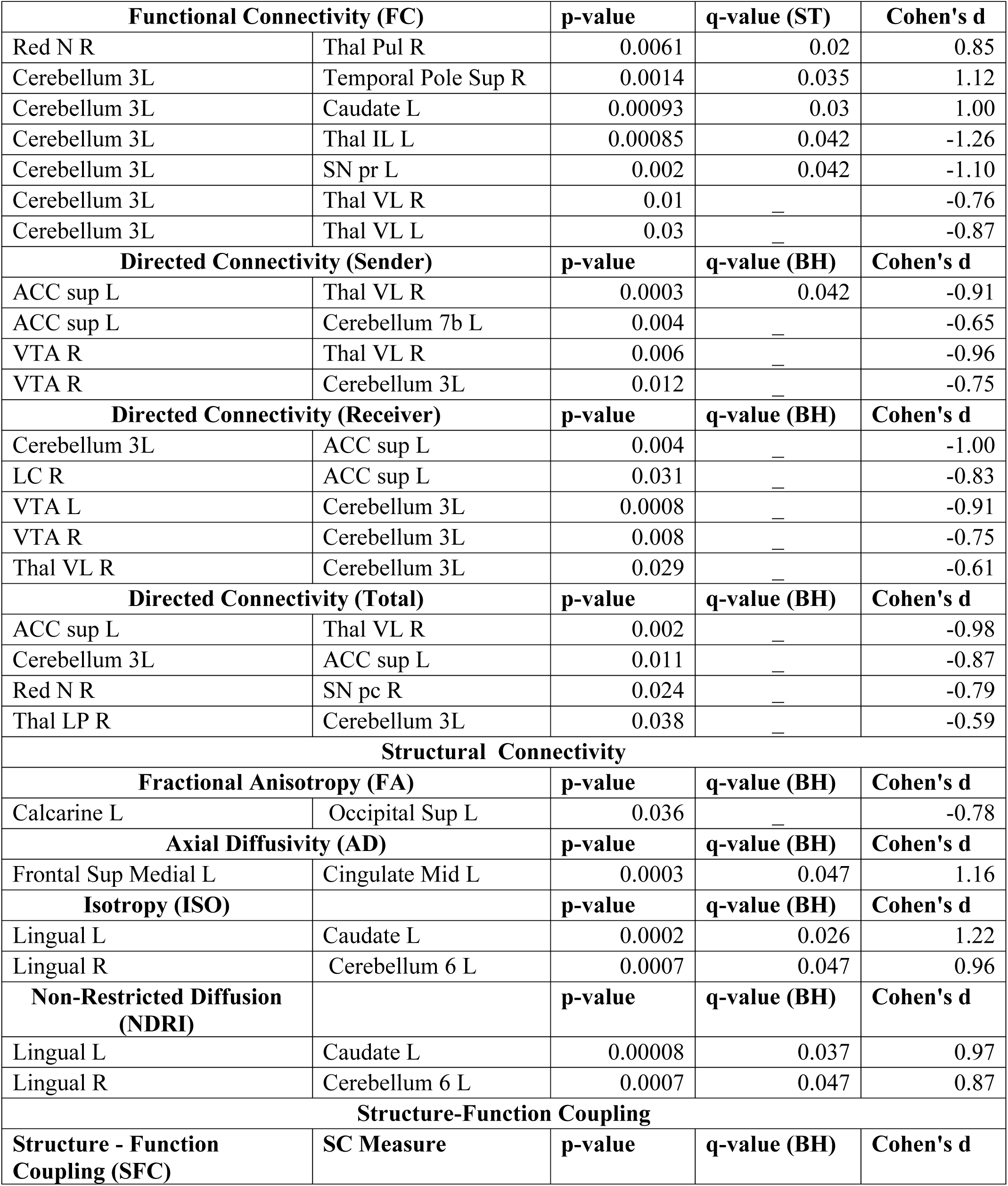
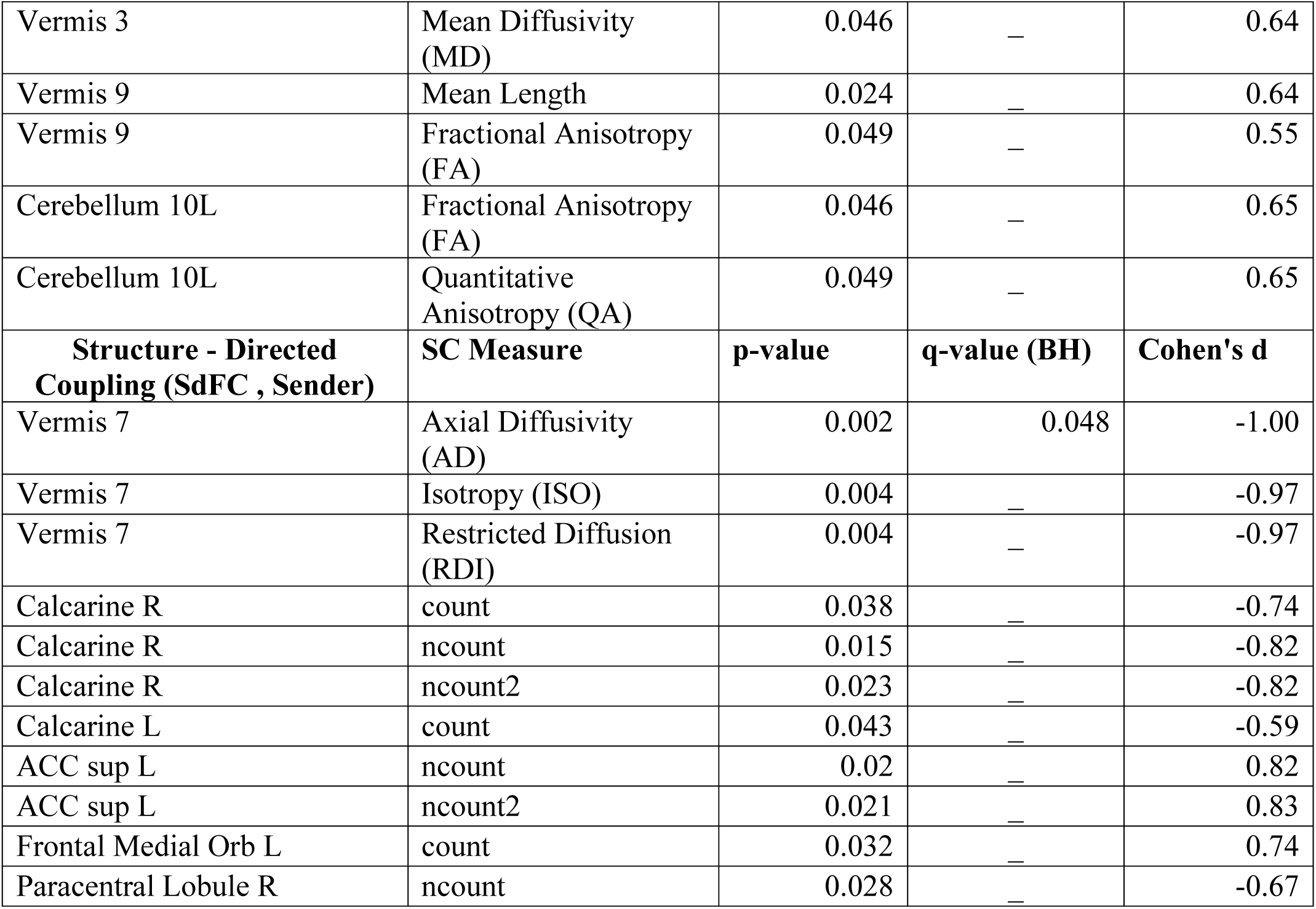
Group-Level Statistics Across Modalities. This figure shows statistical comparisons between AVG players and non-gamers across connectivity measures, including structural, functional, and structure-function coupling. Cohen’s d values indicate effect size, with negative values favoring non-gamers and positive values favoring gamers. Storey-Tibshirani method (ST) and Benjamini-Hochberg (BH) corrections were applied to control for multiple comparisons.

## Author Contributions

**Kyle Cahill:** Conceptualization, Methodology, Software, Formal analysis, Writing - original draft, review & editing. **Mukesh Dhamala:** Conceptualization, Methodology, Software, Supervision, Funding acquisition, Writing - review & editing.

## Data Availability Statement

All data that support the findings of the study as well as the custom analysis scripts can be found in OSF (*link will be public after publication*)

## Ethics Statement

This research was conducted in accordance with all relevant ethical guidelines, including informed consent procedures, participant confidentiality, and adherence to the principles outlined by the Institutional Review Board at Georgia State University. All participants provided written informed consent prior to data collection, and their identities were protected throughout the study.

## Acknowledgments

This work was funded by two internal grant awards of Brains and Behavior Program & the Center for Advanced Brain Imaging to M.D. Bhim Adhikari for providing fMRI preprocessing pipeline. Tim Jordan for initial data collection.

